# Testis-Specific MAGEA3/6 Regulate Glycolysis to Promote Metabolic Flexibility and Reveal Therapeutic Vulnerabilities in Pancreatic Cancer Cells

**DOI:** 10.64898/2026.10.20.760599

**Authors:** Sima Tozandehjani, Vesna Jurjevič, Barbara Breznik Vittori, Juan Sebastian Solano-Gutierrez, Rebecca R. Florke Gee, Denis Štepihar, Maria Camila Hoyos Sanchez, Sara Uhan, Jerneja Koren, Lara Snoj, Emily Ballard, Jesenia M. Perez, Menglin Jiang, Miloš Vittori, Patrick Ryan Potts, Klementina Fon Tacer

## Abstract

Pancreatic ductal adenocarcinoma (PDAC) is an aggressive tumor characterized by metabolic plasticity that enables cancer cells to survive nutrient deprivation and therapeutic stress. Melanoma-associated antigens A3 and A6 (MAGEA3/6), members of the cancer-testis antigen family with a previously identified protective role in spermatogenesis during nutritional stress, have been implicated in cancer progression and metabolic regulation. However, their contribution to metabolic adaptation in pancreatic cancer remains poorly understood. Here, we investigated the molecular mechanisms underlying MAGEA3/6-mediated resistance to glycolytic inhibition using complementary gain- and loss-of-function models, pharmacological inhibition of metabolic pathways, cellular metabolic assays, and integrated transcriptomic, proteomic, metabolomic, and lipidomic analyses. MAGEA3/6 expression enhanced resistance to 2-deoxy-D-glucose (2-DG), with MAGEA6-expressing MIA PaCa-2 cells exhibiting a greater than 2.5-fold increase in the half-maximal inhibitory concentration (IC50). Conversely, MAGEA3/6 depletion reduced clonogenic growth and migration and increased sensitivity to 2-DG. Resistance persisted despite pharmacological inhibition of mTORC1, the pentose phosphate pathway, or glutaminase, suggesting metabolic adaptations beyond these pathways. Unexpectedly, MAGEA6 expression was associated with increased rather than decreased AMPKα1 protein abundance, contrasting with previously reported MAGEA3/6-mediated AMPK degradation and suggesting context-dependent regulation of this master regulator of cellular metabolism. MAGEA6 also reduced hexokinase HK1 and HK2 abundance while increasing several downstream glycolytic enzymes, revealing discordant regulation of glycolysis. Seahorse analyses and integrated multi-omics profiling identified extensive remodeling of mitochondrial metabolism, glutamine-aspartate metabolism, phospholipid composition, lipid storage, and redox homeostasis. Interestingly, despite enhanced resistance to glycolytic inhibition, MAGEA6-expressing cells exhibited increased sensitivity to hydrogen peroxide-induced oxidative stress, revealing a potential metabolic vulnerability. Given that MAGEA3/6 are physiologically expressed primarily in the male germline during early spermatogenesis, a process highly sensitive to metabolic and oxidative stress, these findings also raise important questions about their physiological roles in protecting developing germ cells and maintaining metabolic and redox homeostasis. Together, our findings recognize MAGEA3/6 as regulators of metabolic plasticity in PDAC and highlight how understanding their molecular functions in both spermatogenesis and cancer may uncover previously unrecognized metabolic vulnerabilities and potential therapeutic opportunities.

## INTRODUCTION

Pancreatic ductal adenocarcinoma (PDAC) is one of the deadliest malignancies, with a 5-year survival rate of approximately 13%, largely due to late diagnosis, aggressive progression, early metastasis, and limited treatment options [1–6]. Despite advances in PDAC biology and therapeutic strategies, including surgery, chemotherapy, radiotherapy, and immunotherapy, poor survival and therapeutic resistance remain major clinical challenges. Despite advances in PDAC treatment, poor survival and therapeutic resistance remain major challenges. Emerging targeted therapies, particularly the RAS inhibitor daraxonrasib, have shown promising results, nearly doubling median overall survival compared with standard chemotherapy in previously treated metastatic PDAC [7]. However, the five-year survival rate remains low, highlighting the urgent need to elucidate novel molecular mechanisms underlying PDAC progression and therapeutic resistance, identify new biomarkers for early detection, patient stratification, and prediction of treatment response, and develop more effective targeted therapies [8].

Immunotherapy has transformed treatment outcomes in several types of cancers, but its efficacy in PDAC remains limited [9, 10]. Among therapeutic targets, melanoma-associated antigens (MAGEs), particularly the MAGEA subfamily, have attracted considerable attention due to their restricted expression in the male germline and frequent aberrant activation in cancers, where they are associated with tumor aggressiveness and poor prognosis [11–16]. Since the discovery of MAGEA1 as the first human tumor antigen in 1991 [17], their therapeutic potential has been highlighted by the FDA approval of the first MAGEA4-directed T-cell receptor therapy in 2024 [18]. Recent studies have also implicated MAGEA proteins in PDAC progression and chemoresistance [19]. Especially, MAGEA3 and MAGEA6, which share 96% protein sequence identity (hereafter referred to as MAGEA3/6), are expressed in PDAC and associated with poor clinical outcomes [16, 20–24]. However, their molecular functions and contributions to PDAC progression remain poorly understood.

Insights into their potential roles in cancer have also emerged from studies of their physiological and molecular functions. *MAGEA* genes belong to the cancer–testis antigen family and are normally expressed predominantly in the testis but aberrantly activated in a wide range of cancers [13–15, 25–27]. We have focused on understanding the long-overlooked physiological functions of MAGEA proteins to get new insight into their oncogeneic roles. Our extensive expression profiling revealed that MAGEA genes are expressed in premeiotic male germ cells, just before their entry into meiosis and migration across the blood–testis barrier, a pattern conserved between humans and mice [14].

Intriguingly, under physiological conditions, *MAGEA* genes are predominantly expressed in differentiating spermatogonia approaching meiotic entry, a developmental stage characterized by extensive metabolic reprogramming and increased demand for metabolic plasticity [28, 29]. Our previous studies demonstrated that although *Magea*-deficient mice remain fertile under standard conditions, prolonged food restriction results in more pronounced spermatogenic defects and reduced sperm counts compared with wild-type mice [28]. These findings suggest that MAGEA proteins may have evolved to protect germ cells during periods of nutritional stress, a function potentially hijacked by cancer cells to support their metabolic adaptation and survival. Supporting this hypothesis, our preliminary studies demonstrated that MAGEA expression protects cancer cells against glycolytic inhibition by 2-deoxy-D-glucose (2DG) [28]. However, the molecular mechanisms underlying MAGEA-mediated metabolic adaptation remain unanswered.

Understanding of the molecular functions of MAGEA proteins have largely emerged from cancer studies. MAGEA proteins interact with TRIM28 (KAP1/TIF1β), a multifunctional transcriptional coregulator and E3 ubiquitin ligase, to regulate transcription and promote ubiquitination of specific substrates [13]. Interestingly, beyond p53, several identified targets are key metabolic regulators, including AMP-activated protein kinase (AMPK) and fructose-1,6-bisphosphatase 1 (FBP1) [30–34]. MAGEA3/6-mediated AMPK degradation suppresses autophagy and promotes cancer cell survival, while FBP1 downregulation enhances glucose consumption and lactate production, promoting the Warburg effect [20, 22, 31, 35–42]. In all, these findings suggest that MAGEA proteins contribute to metabolic reprogramming, an established hallmark of cancer and an emerging therapeutic vulnerability.

Rapidly proliferating PDAC cells rely on oncogene-driven metabolic reprogramming to meet their increased energetic and biosynthetic demands while adapting to the nutrient-deprived and hypoxic tumor microenvironment [43, 44]. Glucose metabolic reprogramming is one of the hallmarks of pancreatic cancer, with oncogenic KRAS promoting glycolysis and redirecting glucose intermediates toward biosynthetic pathways, including the hexosamine biosynthesis pathway and non-oxidative pentose phosphate pathway. In addition to glucose, glutamine is a major carbon source that supports TCA cycle activity, biosynthesis, and redox homeostasis, with PDAC cells exhibiting a strong dependence on glutamine for sustained growth and proliferation [45]. Lipid metabolism is another critical component of PDAC metabolic reprogramming, providing alternative energy sources under metabolic stress while supporting cellular signaling, membrane remodeling, and tumor cell survival and proliferation [46]. Collectively, these metabolic adaptations enable cancer cells to maintain redox homeostasis and sustain proliferation under nutrient-limited and oxidative stress conditions. However, despite their frequent expression in cancer and the similarly unique metabolic demands of spermatogenesis, how cancer–testis genes contribute to metabolic adaptation remains largely unknown.

Targeting cancer metabolism has achieved clinical success in selected malignancies, particularly those with specific metabolic vulnerabilities. However, targeting glycolysis and glutamine metabolism in solid tumors, including PDAC, has been less successful, potentially due to the metabolic plasticity of cancer cells and their ability to engage alternative pathways to survive metabolic stress [47, 48]. For example, 2-deoxy-D-glucose (2-DG) initially emerged as a promising glycolytic inhibitor, but its clinical efficacy has been limited, partly due to adaptive resistance mechanisms that are only beginning to be understood. Nevertheless, 2-DG remains a valuable tool for investigating metabolic dependencies and adaptive responses [49], an approach we exploited in this study.

Here, we aimed to elucidate the molecular mechanisms underlying MAGEA3/6-mediated resistance to 2-DG in highly glycolytic PDAC cells and uncover previously unrecognized roles of MAGEA3/6 in metabolic reprogramming. Using complementary gain- and loss-of-function models, we investigated responses to glycolytic inhibition, pentose phosphate pathway inhibition, and glutamine deprivation by integrating functional assays with transcriptomic, proteomic, metabolomic, and lipidomic analyses. Our findings reveal a previously unrecognized mechanism of metabolic adaptation characterized by discordant regulation of glycolytic enzymes, with reduced hexokinase abundance despite increased levels of other glycolytic proteins. These changes are accompanied by enhanced cellular bioenergetics and altered lipid metabolism, apparently independent of the previously described AMPK/mTOR signaling axis. Collectively, our findings identify MAGEA3/6 as a regulator of metabolic plasticity that promotes PDAC cell survival under metabolic stress, reveal previously unrecognized molecular targets of MAGEA3/6, and highlight potential therapeutic vulnerabilities.

## MATERIALS AND METHODS

### Reagents and Materials

#### Chemicals, inhibitors, and supplements

2-Deoxy-D-glucose (2-DG; Cat. No. D6134-5G), 6-aminonicotinamide (6-AN; Cat. No. A68203), dimethyl sulfoxide (DMSO; Cat. No. D2650-100ML), 2-mercaptoethanol (Cat. No. M3148-25ML), poly-D-lysine hydrobromide (Cat. No. P7280-5MG), puromycin dihydrochloride (Cat. No. P8833-100MG), doxycycline hydrochloride (Cat. No. D3447-500MG), rapamycin (Cat. No. R0395-1MG), DAPI (Cat. No. D9542-5MG), bis-2-(5-phenylacetamido-1,3,4-thiadiazol-2-yl)ethyl sulfide (BPTES; Cat. No. SML0601-5MG), doxorubicin hydrochloride (Cat. No. D1515-10MG), reduced L-glutathione (Cat. No. G6529-5G), horse serum (Cat. No. H1270-500ML), BioXtra L-glutamine solution (200 mM; Cat. No. G7513-100ML), SIGMAFAST Protease Inhibitor Cocktail Tablets, EDTA-free (Cat. No. S8830-20TAB), and Protease and Phosphatase Inhibitor Cocktail (Cat. No. PPC1010-1ML) were purchased from Sigma-Aldrich (St. Louis, MO, USA). Bovine serum albumin (BSA), Fraction V (Cat. No. 126609-10GM), was obtained from MilliporeSigma (Burlington, MA, USA). PhosSTOP Phosphatase Inhibitor (Cat. No. 04906845001) was obtained from Roche (Indianapolis, IN, USA).

#### Cell culture media, transfection reagents, and molecular biology reagents

DMEM high glucose (Cat. No. 11965092), glucose-free DMEM (Cat. No. 11966025), glutamine-free high-glucose DMEM (Cat. No. 11960044), RPMI 1640 medium (Cat. No. 11875093), penicillin–streptomycin (Cat. No. 15140122), trypsin-EDTA (Cat. No. 25200056), sodium pyruvate (Cat. No. 11360070), Opti-MEM I Reduced Serum Medium (Cat. No. 11058021), and DPBS without calcium and magnesium (Cat. No. 14190144) were obtained from Gibco (Grand Island, NY, USA). Lipofectamine 3000 (Cat. No. L3000015), Lipofectamine RNAiMAX (Cat. No. 13778150), Trypan Blue Stain (Cat. No. T10282), Oligo(dT)20 Primer (Cat. No. 18418020), and SuperScript IV Reverse Transcriptase (Cat. No. 18090050) were obtained from Invitrogen (Carlsbad, CA, USA). PowerUp SYBR Green Master Mix (Cat. No. A25742), PowerTrack SYBR Green Master Mix (Cat. No. A46109), and High-Capacity cDNA Reverse Transcription Kit (Cat. No. 4368814) were obtained from Applied Biosystems (Foster City, CA, USA). The Quick-RNA Miniprep Plus Kit was obtained from Zymo Research (Irvine, CA, USA).

#### Fluorescence imaging, protein analysis, and metabolic assays

Hoechst 33342 (Cat. No. H3570), MitoTracker Red CMXRos (Cat. No. M7512), and HCS LipidTOX Deep Red Neutral Lipid Stain (Cat. No. H34477) were obtained from Molecular Probes (Eugene, OR, USA). IRDye 800CW 2-DG Optical Probe (Cat. No. 926-08946) was obtained from LI-COR Biosciences (Lincoln, NE, USA). Pierce RIPA Lysis and Extraction Buffer (Cat. No. 89901), Ponceau S Staining Solution (Cat. No. A40000279), and Pierce Protein A Agarose (Cat. No. 20333) were obtained from Thermo Scientific (Rockford, IL, USA). Anti-rabbit IgG (H+L), HRP-conjugated secondary antibody (Cat. No. W4011), was obtained from Promega Corporation (Madison, WI, USA). Alamar Blue Reagent (Cat. No. BUF012B), SilentFect (Cat. No. 1703361), and Trans-Blot Turbo 5× Transfer Buffer (Cat. No. 10026938) were obtained from Bio-Rad Laboratories (Hercules, CA, USA). Seahorse XF Calibrant Solution (Cat. No. 100840-000), Seahorse XF DMEM Medium, pH 7.4 (Cat. No. 103575-100), Seahorse XF Long Chain Fatty Acid Oxidation Stress Test Kit (Cat. No. 103672-100), XF24 Extracellular Flux Assay Kit (Lot No. B05423), Seahorse XF24 Cell Culture Microplates (Lot No. 09918), and Seahorse XF Real-Time ATP Rate Assay Kit (Cat. No. 103592-100) were obtained from Agilent Technologies (Cedar Creek, TX, USA).

#### General laboratory consumables and sample preparation reagents

PBS without calcium and magnesium (Cat. No. 21-040-CV), Costar 96-well plates (Ref. 3596), Transwell permeable supports (Ref. 3412), Costar multiwell plates, 10-cm tissue-culture dishes (Ref. 353003), and 5-mL and 10-mL serological pipettes (Refs. 357543 and 357551) were obtained from Corning Incorporated (Corning, NY, USA). Paraformaldehyde solution, 4% in PBS (Cat. No. J19943.K2), was obtained from Thermo Fisher Scientific (Waltham, MA, USA). Richard-Allan Scientific Mounting Medium (Cat. No. 84000-112) and sterile 200-µL low-retention filtered pipette tips (Cat. No. 76322-150) were obtained from VWR International (Radnor, PA, USA). Sterile 5-mL serological pipettes (Cat. No. 86.1253.025) were obtained from Sarstedt (Newton, NC, USA). Electron microscopy-grade glutaraldehyde (Cat. No. 18426) was obtained from Ted Pella, Inc. (Redding, CA, USA).

### Cell Lines and Culture Conditions

Human pancreatic ductal adenocarcinoma (PDAC) cell lines MIA PaCa-2, BxPC-3, DAN-G, PANC-1, and PANC-1-Luc2, as well as human embryonic kidney HEK293 cells, were used in this study. MIA PaCa-2 cells engineered to overexpress *MAGEA6*, along with GFP-expressing control cells, were generously provided by Dr. Kenneth L. Scott [14, 37]. MIA PaCa-2 cells were maintained in high-glucose Dulbecco’s Modified Eagle Medium (DMEM) supplemented with 10% fetal bovine serum (FBS), 2.5% horse serum, and 1× antibiotic–antimycotic solution. BxPC-3 cells (ATCC, CRL-1687) were maintained in RPMI-1640 medium supplemented with 10% FBS and 1× antibiotic–antimycotic solution. DAN-G cells (Cytion, Cat. No. 300162, Sioux Falls, SD, USA) were cultured in RPMI-1640 medium containing 2.1 mM stable L-glutamine and 2.0 g/L sodium bicarbonate, supplemented with 10% FBS and 1× antibiotic–antimycotic solution. PANC-1 (ATCC, CRL-1469) and PANC-1-Luc2 (ATCC, CRL-1469-LUC2) cells, used for knockdown and overexpression studies, respectively, were maintained in DMEM (ATCC, Cat. No. 30-2002) supplemented with 10% FBS and 1× antibiotic–antimycotic solution. PANC-1-Luc2 cells were additionally maintained under selection with 8 µg/mL blasticidin. HEK293 cells, originally obtained from GenTarget Inc. (CA, USA; Cat. No. SC062-LG) as a putative AsPC-1 derivative stably expressing luciferase and GFP with puromycin resistance, were subsequently identified as HEK293 by cell-line authentication. These cells were maintained in RPMI-1640 medium containing HEPES and high glucose, supplemented with 10% FBS and 1× antibiotic–antimycotic solution. All cell lines were cultured at 37 °C in a humidified atmosphere containing 5% CO₂. Cell numbers and viability were determined using a hemocytometer or a Countess II Automated Cell Counter (Invitrogen, Thermo Fisher Scientific, Waltham, MA, USA) with trypan blue exclusion.

### Cell Line Authentication

Cell line identity was confirmed by short tandem repeat (STR) profiling performed at the Pediatric Cancer Research Center, Texas Tech University Health Sciences Center (TTUHSC), using the GenePrint 10 PCR Amplification Kit (Promega) according to the manufacturer’s instructions. For authentication, cells were cultured to approximately 95% confluence, washed with phosphate-buffered saline (PBS), and detached by trypsinization for 5–15 min, depending on the cell line. For MIA PaCa-2 cells, culture medium containing floating cells was collected before washing and retained to minimize cell loss. Detached cells were collected in complete medium and centrifuged at 300 × g at room temperature. Cell pellets were resuspended in PBS, and cells were counted. Approximately 2–6 × 10⁶ cells were collected, washed again with PBS, pelleted by centrifugation, and snap-frozen in liquid nitrogen. Samples were subsequently analyzed at the Pediatric Cancer Research Center, TTUHSC. STR profiles were compared with established reference profiles to verify cell line identity.

### Genomic DNA Extraction and Mycoplasma Detection

Genomic DNA was extracted from cell culture samples using the G-spin™ Total DNA Extraction Mini Kit according to the manufacturer’s instructions. Briefly, 200 µL of cell culture material was mixed with 200 µL of Buffer ML1, 20 µL of proteinase K, and 5 µL of RNase A, followed by incubation at 56 °C for 10 min. Following cell lysis, Buffer ML2 and absolute ethanol were added, and the lysate was transferred to a spin column. The column was sequentially washed with MWA and MWB buffers, and genomic DNA was eluted in 50 µL of Buffer ME by centrifugation. Purified genomic DNA was subsequently used for PCR-based mycoplasma detection using the e-Myco™ Mycoplasma PCR Detection Kit (Boca Scientific; Fisher Scientific, Cat. No. 501685641) according to the manufacturer’s instructions. All cell lines tested negative for mycoplasma contamination.

### Generation of Stable Overexpression Cell Lines

Stable overexpression cell lines were generated by transfecting MIA PaCa-2 cells with Lipofectamine 3000 (Invitrogen, Thermo Fisher Scientific, Waltham, MA, USA) according to the manufacturer’s instructions. Briefly, 2 × 10⁴ cells were seeded per well in 6-well plates and cultured for 24 h before transfection. Cells were transfected with 1 µg of plasmid DNA per well, including pIRES-puro-TAP-MAGEA3, pIRES-puro-TAP-MAGEA6, or the corresponding control plasmids pIRES-puro-mCherry, pIRES-puro-TAP-Tag, and pIRES-puro-FLAG [30]. At 48 h post-transfection, cells were trypsinized and replated into 10-cm culture dishes. Following an additional 24 h, selection was initiated with 1 µg/mL puromycin. After 2–4 weeks of selection, stable expression of mCherry, FLAG, and TAP-tagged proteins was confirmed by Western blotting. Cells were subsequently maintained in medium containing 1 µg/mL puromycin for several months to enrich for stably transfected cells.

### siRNA-Mediated Knockdown of MAGEA3/6

Transient knockdown of *MAGEA3/6* was performed in BxPC-3 cells using SilentFect (Bio-Rad Laboratories, Hercules, CA, USA) and in MIA PaCa-2, PANC-1, and DAN-G cells using Lipofectamine RNAiMAX (Invitrogen, Thermo Fisher Scientific, Waltham, MA, USA). Briefly, BxPC-3 (1 × 10⁶ cells) and DAN-G (3 × 10⁶ cells) were seeded in 10-cm culture dishes and allowed to adhere for 24 h. Before transfection, complete growth medium was replaced with Opti-MEM Reduced Serum Medium for 2 h. Cells were subsequently transfected with small interfering RNAs (siRNAs) targeting ubiquitin B (UBB; oligo Nos. 3030837381-000300 and 3030837381-000310), reverse transcriptase-like sequence (siRev-L1; oligo No. WD04812532), LonRF (oligo Nos. 3012482021-000160 and 3012482021-000170), or MAGEA3/6 (si-panMAGEA3/6; oligo No. WD08299911). All siRNAs were obtained from Sigma-Aldrich (St. Louis, MO, USA) and transfected at a final concentration of 33.3 nM according to the manufacturer’s instructions. After 24 h, the transfection medium was replaced with fresh complete growth medium, and cells were maintained for a total of 72 h post-transfection before harvesting for downstream analyses. PANC-1 cells were transfected with the same siRNAs using Lipofectamine RNAiMAX according to the manufacturer’s protocol, without prior serum starvation.

### Cell Viability Assay

MIA PaCa-2 cells were seeded in 96-well plates at a density of 1 × 10⁴ cells per well in complete or glutamine-free culture medium. After 24 h, one plate maintained in complete medium was analyzed to establish baseline cell viability. Cells were subsequently treated with increasing concentrations of 2-deoxy-D-glucose (2-DG) in complete or glutamine-free medium, BPTES in complete or glucose-free medium, or 3-bromopyruvate (3-BP) in complete medium, as indicated in the corresponding figure legends. Cell viability was assessed at 72 and 120 h post-treatment using the AlamarBlue assay. Briefly, 90 µL of culture medium was removed from each well, followed by the addition of 10 µL of AlamarBlue reagent. Plates were incubated for 4 h at 37 °C, and fluorescence was measured using a Cytation 5 Cell Imaging Multimode Reader (BioTek Instruments, Winooski, VT, USA). Fluorescence values were normalized to untreated controls and expressed as relative cell viability. Experiments were performed in three independent biological replicates. Dose–response curves were fitted using nonlinear regression in GraphPad Prism (GraphPad Software, Boston, MA, USA) to determine half-maximal inhibitory concentrations (IC50). The concentrations tested are specified in the corresponding figure legends.

### Treatment with 2-Deoxy-D-glucose (2-DG)

MIA PaCa-2-GFP and MIA PaCa-2-MAGEA3/6 cells were seeded at a density of 1 × 10⁶ cells per 6-cm dish and cultured for 24 h to approximately 90% confluence. Cells were subjected to five experimental conditions: (1) complete medium (untreated control), (2) complete medium containing 3 mM 2-DG for 24 h, (3) complete medium containing 3 mM 2-DG for 48 h, (4) glutamine-free medium, and (5) glutamine-free medium containing 3 mM 2-DG for 24 h. For the combined treatment, glutamine deprivation and 2-DG exposure were initiated simultaneously. Each condition was prepared in technical duplicate, and experiments were independently repeated three times. Following treatment, floating cells were collected, and adherent cells were washed with phosphate-buffered saline (PBS) and detached by trypsinization for 10 min. Both cell fractions were combined and centrifuged at 300 × *g* for 5 min. The resulting cell pellets were collected for downstream transcriptomic, proteomic, metabolomic, and lipidomic analyses, as described below.

### Assessment of Cellular 2-DG Uptake

Cellular uptake of 2-deoxy-D-glucose (2-DG) was assessed using the IRDye 800CW 2-DG Optical Probe (LI-COR Biosciences, Lincoln, NE, USA). MIA PaCa-2-GFP and MIA PaCa-2-MAGEA6 cells were seeded at a density of 2.5 × 10⁵ cells per well in 24-well plates and cultured for 24 h. Complete culture medium was subsequently replaced with glucose-free medium, and cells were incubated with 250 µL of IRDye 800CW 2-DG probe (5 µM) for 6 h. Following incubation, cells were washed twice with phosphate-buffered saline (PBS) to remove unbound probe and fixed with 4% paraformaldehyde (PFA) for 20 min at room temperature [50]. Nuclei were counterstained with DAPI, and fluorescence images were acquired using a Leica STELLARIS 8 FALCON STED super-resolution confocal microscope (Leica Microsystems). IRDye 800CW 2-DG fluorescence was detected using excitation wavelengths of approximately 774–780 nm and an emission detection range of 785–850 nm. GFP and DAPI fluorescence were detected using emission ranges of approximately 500–550 nm and 420–480 nm, respectively.

### RNA Extraction and Purification

Total RNA was extracted using the Quick-RNA Miniprep Plus Kit (Zymo Research, Irvine, CA, USA) according to the manufacturer’s instructions, with minor modifications to improve RNA purity and concentration. Briefly, cell lysates were subjected to spin-column purification with on-column DNase I digestion, extended to 15 min to minimize genomic DNA contamination. RNA was eluted in a reduced volume of DNase/RNase-free water to increase RNA concentration (15 µL). RNA yield and purity were assessed using a NanoDrop 2000 spectrophotometer (Thermo Scientific, Waltham, MA, USA), and purified RNA was used for downstream analyses.

### RNA Sequencing (RNA-seq) and Differential Gene Expression Analysis

MIA PaCa-2-GFP and MIA PaCa-2-MAGEA3/6 cells were cultured in complete or glutamine-free medium and treated with 2-deoxy-D-glucose (2-DG), as described above. Total RNA was extracted and its concentration and purity assessed as described previously. For each sample, 40 ng of total RNA in 1 µL of nuclease-free water was submitted to Novogene (Sacramento, CA, USA) for library preparation and sequencing. RNA-seq libraries were prepared by poly(A) mRNA enrichment using oligo(dT) magnetic beads, followed by mRNA fragmentation, first-strand cDNA synthesis using random hexamer primers, and second-strand cDNA synthesis. Libraries were sequenced on an Illumina NovaSeq platform using 150-bp paired-end (PE150) sequencing with sequencing-by-synthesis (SBS) chemistry.

Raw sequencing data were processed for base calling and converted into FASTQ files, and sequencing quality was assessed using FastQC. Sequencing reads were aligned to the human reference genome (GRCh38/hg38) using HISAT2 (v2.0.5) [51]. Gene expression was quantified using raw gene-level read counts and fragments per kilobase of transcript per million mapped reads (FPKM). Principal component analysis (PCA) was performed using FPKM values to assess intergroup differences and intragroup sample clustering. Differential gene expression analysis was performed using the DESeq2 R package (v1.20.0) [52], with normalization of raw gene counts to account for differences in sequencing depth. *P*-values were adjusted for multiple comparisons using the Benjamini–Hochberg false discovery rate (FDR) method. Genes with an adjusted *p*-value (padj) ≤ 0.05 and an absolute log₂ fold change ≥ 1 were considered differentially expressed.

### cDNA Synthesis and Quantitative Real-Time PCR (RT-qPCR)

For cDNA synthesis, 4 µg of total RNA was treated with DNase to remove residual genomic DNA and reverse transcribed using random primers in a 40-µL reaction. Reverse transcription was performed at 25 °C for 10 min, 37 °C for 2 h, and 85 °C for 5 min, followed by a hold at 4 °C. The resulting cDNA was diluted with nuclease-free water and used for quantitative real-time PCR (qPCR). qPCR was performed using SYBR Green chemistry in 10-µL reactions containing SYBR Green PCR master mix, gene-specific primers, and diluted cDNA. Amplification was carried out for 40 cycles at 95 °C for 15 s and 60 °C for 60 s, with fluorescence acquisition during the extension step [53]. Primer sequences and experimental procedures were as previously reported [14].

### Metabolite, Lipid, and Protein Extraction

#### Metabolite and lipid extraction

Approximately 3 × 10⁶ cells were washed, resuspended in 500 µL of phosphate-buffered saline (PBS), snap-frozen, and submitted to the Center for Biotechnology and Genomics (CBG) at Texas Tech University (Lubbock, TX, USA) for analysis. Cell suspensions were centrifuged at 5,000 rpm for 15 min at 4 °C, and the resulting pellets were resuspended in 100 µL of water following PBS removal. EQUISPLASH™ LIPIDOMIX™ Quantitative Mass Spec Internal Standard (Cat. No. 330731) and L-phenyl-d5-alanine-2,3,3-d3 (Cat. No. 490148; Sigma-Aldrich, St. Louis, MO, USA) were added, followed by two ceramic beads (Cat. No. 13114-325; Qiagen, Germantown, MD, USA).

Metabolites and lipids were extracted using dichloromethane (1:2, v/v), followed by bead-mill homogenization (Mini-Beadbeater-96, BioSpec Products, Bartlesville, OK, USA), vortexing, and sonication on ice. Phase separation was achieved by sequential addition of dichloromethane and cold water, followed by homogenization and centrifugation at 10,000 × *g* for 15 min at 4 °C. The upper aqueous phase containing polar metabolites and lower organic phase containing nonpolar lipids were collected separately. Polar extracts were clarified by centrifugation and prepared for liquid chromatography–mass spectrometry (LC-MS), whereas lipid extracts were dried and resuspended in acetonitrile (65:30:5, v/v/v) before LC-MS analysis. The remaining pellets were washed with methanol and stored at −80 °C for subsequent protein extraction. **Protein extraction.** The remaining pellets were resuspended in 100 µL of extraction buffer containing 50 mM ammonium bicarbonate and 2.5% (w/v) sodium deoxycholate (SDC). Samples were vortexed, homogenized using a bead mill at 2,400 Hz for 45 s, and sonicated on ice for 1 h to facilitate protein solubilization. Following centrifugation at 10,000 × *g* for 5 min, protein-containing supernatants were collected for subsequent liquid chromatography–tandem mass spectrometry (LC-MS/MS) analysis.

### Liquid Chromatography–Tandem Mass Spectrometry (LC–MS/MS) Analysis

Multi-omics analyses, including metabolomics, lipidomics, and proteomics, were performed at the Center for Biotechnology and Genomics (CBG), Texas Tech University (Lubbock, TX, USA). **Metabolomics.** Polar metabolites were analyzed using a Vanquish Horizon ultra-high-performance liquid chromatography (UHPLC) system coupled to a Q Exactive HF Orbitrap mass spectrometer (Thermo Fisher Scientific). Chromatographic separation was performed on a Waters ACQUITY UPLC HSS T3 column (2.1 × 100 mm, 1.7 µm) maintained at 55 °C, with a flow rate of 0.4 mL/min and an injection volume of 5 µL. Mobile phases consisted of water (A) and methanol (B), both containing 0.1% formic acid. Mass spectrometry data were acquired in positive and negative electrospray ionization modes using full-scan MS and data-dependent tandem mass spectrometry (ddMS2). Full MS and MS/MS spectra were acquired at resolutions of 120,000 and 30,000, respectively, over an *m/z* range of 250–1200. **Lipidomics.** Nonpolar lipids were analyzed using the same UHPLC–MS system at 55 °C, with a flow rate of 0.35 mL/min and an injection volume of 5 µL. Mobile phases consisted of acetonitrile/water (A) and isopropanol/acetonitrile/water (B), each containing 10 mM ammonium formate and 0.1% formic acid. Data were acquired in positive and negative electrospray ionization modes using full-scan MS and ddMS2, with resolutions of 120,000 and 30,000, respectively, over an *m/z* range of 250–1200. **Proteomics.** Proteomic analyses were performed using an UltiMate 3000 RSLCnano system coupled to a Q Exactive HF Orbitrap mass spectrometer (Thermo Fisher Scientific). Peptides were separated on a Thermo PepMap RSLC C18 column (75 µm × 150 mm, 3 µm) maintained at 50 °C, using a binary mobile phase consisting of 0.1% formic acid in water (A) and 0.1% formic acid in acetonitrile (B). A linear gradient from 2% to 30% B was applied, followed by increases to 50% and 90% B for column washing and subsequent re-equilibration. The flow rate was maintained at 0.30 µL/min, with an injection volume of 2 µL. Mass spectrometry data were acquired in positive electrospray ionization mode using data-dependent acquisition (DDA). Full MS scans were collected at a resolution of 120,000, an automatic gain control (AGC) target of 1 × 10⁶, a maximum injection time of 50 ms, and an *m/z* range of 300–1800. MS/MS spectra were acquired at a resolution of 15,000, an AGC target of 1 × 10⁵, and a maximum injection time of 50 ms, using a normalized collision energy of 27 and an isolation window of 1.6 *m/z*.

### Analysis of Metabolomic, Lipidomic, and Proteomic Data

Metabolomic and lipidomic datasets were analyzed separately using MetaboAnalyst. Technical replicates were averaged for each biological sample, resulting in three independent biological replicates per experimental group. Low-variance features were filtered based on the interquartile range (IQR). Metabolomic data were normalized to the sum of feature intensities, whereas lipidomic data retained the normalization applied during upstream processing. Missing lipid values were imputed using one-fifth of the minimum positive value. Both datasets were log₁₀-transformed and Pareto-scaled before statistical analysis. Principal component analysis (PCA) was performed to evaluate sample clustering and overall variation. Statistical differences were assessed using one-way analysis of variance (ANOVA) and pairwise *t*-tests, followed by correction for multiple testing. Features with a false discovery rate (FDR)-adjusted *P* value < 0.05 were considered statistically significant and further evaluated by fold-change analysis and volcano plots.

Proteomic data were processed and analyzed using ProteoAnalyst. Unannotated proteins and features with more than 70% missing values were excluded, and low-variance features were filtered using a 15% interquartile range (IQR) cutoff. Protein abundances were normalized to the sample median, log₂-transformed, and median-centered. Missing values were imputed using the minimum probability (MinProb) method under a left-censored missing-not-at-random (MNAR) assumption. Differential protein abundance between MAGEA6-expressing and GFP-control MIA PaCa-2 cells under each experimental condition was assessed using the limma framework, which employs linear modeling with empirical Bayes moderation. Proteins with unadjusted *P* < 0.05 were considered nominally significant and included in exploratory downstream analyses. Multiple-testing correction was performed using the Benjamini–Hochberg false discovery rate (FDR) method; however, no proteins met the FDR-adjusted significance threshold of 0.05. Volcano plots were generated using log₂ fold changes (MAGEA6/GFP) and −log₁₀-transformed unadjusted *P* values to visualize nominal differences in protein abundance.

### Seahorse ATP Production Rate Assay

MIA PaCa-2-GFP and MIA PaCa-2-MAGEA6 cells were seeded at a density of 1 × 10⁶ cells per 6-cm dish and cultured for 24 h before treatment with 2-deoxy-D-glucose (2-DG) in complete or glutamine-free medium, as described above. The day before the assay, Seahorse XF24 cell culture microplates (Agilent Technologies, Santa Clara, CA, USA) were coated with 200 µL of poly-D-lysine hydrobromide (Sigma-Aldrich, Cat. No. P-0899) per well for 1 h at room temperature, washed three times with phosphate-buffered saline (PBS), and stored at 4 °C. Sensor cartridges were hydrated overnight in 1 mL of Seahorse XF Calibrant (Cat. No. 100840-000) at 37 °C in a non-CO₂ incubator. On the day of the assay, Seahorse XF assay medium was supplemented with 10 mM glucose, 1 mM pyruvate, and 2 mM glutamine, with glutamine omitted for glutamine-free conditions. Coated microplates were prewarmed at 37 °C for at least 3 h before cell seeding. Following treatment, cells were collected, counted, and seeded into the prewarmed microplates in assay medium. After 1 h of attachment, additional assay medium was gently added, and plates were incubated at 37 °C in a non-CO₂ incubator for 45–60 min before measurement. Cellular ATP production rates were determined using the Seahorse XF Real-Time ATP Rate Assay Kit (Cat. No. 103592-100) and Seahorse XFe24 Extracellular Flux Analyzer (Agilent Technologies). Oligomycin (1.5 µM final concentration) and rotenone/antimycin A (0.5 µM final concentration) were sequentially injected through ports A and B, respectively, according to the manufacturer’s instructions. Following the assay, cells were fixed with 4% paraformaldehyde (PFA) for 10 min at room temperature, washed three times with PBS, stained with Hoechst 33342, and imaged using a Cytation 5 Cell Imaging Multimode Reader (BioTek Instruments, Winooski, VT, USA). Nuclear counts were used to normalize Seahorse measurements to cell number, and glycolytic, mitochondrial, and total ATP production rates were calculated using the manufacturer’s analysis software.

### Mitochondrial and Lipid Staining and Confocal Imaging of MIA PaCa-2 Cells

Mitochondria were stained using MitoTracker Red CMXRos (Molecular Probes, Eugene, OR, USA). Lyophilized MitoTracker was reconstituted in anhydrous dimethyl sulfoxide (DMSO) to prepare a 1 mM stock solution, which was diluted in complete growth medium to final concentrations of 50 or 100 nM. MIA PaCa-2 cells (2 × 10⁵) were seeded onto glass coverslips and cultured for 24 h. The medium was subsequently replaced with prewarmed (37 °C) MitoTracker staining solution, and cells were incubated for 30 min at 37 °C in 5% CO₂. Following incubation, cells were washed three times with phosphate-buffered saline (PBS), fixed with 4% paraformaldehyde (PFA) for 10 min at room temperature, and washed three additional times with PBS. For neutral lipid staining, fixed cells were incubated with HCS LipidTOX Deep Red Neutral Lipid Stain (Molecular Probes, Eugene, OR, USA), diluted 1:1,000 (1×) in culture medium, for 30 min at room temperature. Nuclei were counterstained with DAPI, and cells were imaged by confocal microscopy. Fluorescence images were acquired using a Leica STELLARIS 8 FALCON STED super-resolution confocal microscope (TTUHSC School of Pharmacy Imaging Core, Amarillo, TX, USA). MitoTracker Red CMXRos was excited at approximately 579 nm, with an emission maximum of 599 nm, whereas HCS LipidTOX Deep Red Neutral Lipid Stain was excited at approximately 637 nm, with an emission maximum of 655 nm.

Fluorescence images were analyzed using Fiji/ImageJ [54]. MitoTracker fluorescence was quantified by measuring mean fluorescence intensity (MFI) within cellular regions and subtracting background MFI determined from cell-free regions. The signal-to-background ratio (SBR) was calculated as cellular MFI divided by background MFI. Both parameters were evaluated, with corrected MFI used as the primary measure for visualization and statistical analysis. Fluorescence intensity was expressed in arbitrary units (AU) [55]. Lipid droplets were quantified using Fiji/ImageJ as previously described [56, 57]. LipidTOX Deep Red fluorescence images were thresholded, and lipid droplets were counted using the Analyze Particles function. The total number of lipid droplets was normalized to the number of DAPI-positive nuclei in each image to determine lipid droplets per cell. Statistical analyses and data visualization were performed using GraphPad Prism version 10 (GraphPad Software, Boston, MA, USA).

### Transmission Electron Microscopy (TEM) Sample Preparation and Image Analysis

MIA PaCa-2 cells were cultured and treated as described in the *Treatment with 2-Deoxy-D-glucose (2-DG)* section. At the indicated time points, cells were washed, detached by trypsinization, collected in 5 mL of complete medium, and centrifuged to obtain cell pellets. Following removal of the supernatant, pellets were resuspended in 1.5 mL of phosphate-buffered saline (PBS), transferred to microcentrifuge tubes, and centrifuged again. Cells were fixed in 2.5% electron microscopy-grade glutaraldehyde prepared in 1× PBS by combining 8 mL of ultrapure water, 1 mL of 10× PBS, and 1 mL of 25% glutaraldehyde. Cell pellets were resuspended in prewarmed (37 °C) fixative, incubated for 1 h at room temperature, and stored at 4 °C until further processing for TEM analysis. TEM images were analyzed using ImageJ (version 2.16.0/1.54p) [58]. Images were calibrated using their corresponding scale bars. For each cell, total cell and nuclear areas were measured, and the nuclear area was subtracted from the total cell area to determine the cytoplasmic area. Mitochondria and lipid droplets were counted within the cytoplasmic region, and their respective densities were calculated as the number of organelles per unit cytoplasmic area (µm²). The total area occupied by lipid droplets was measured, and the lipid droplet area fraction was calculated as the ratio of total lipid droplet area to cytoplasmic area, expressed as a percentage. Data were analyzed and visualized using GraphPad Prism version 10 (GraphPad Software, Boston, MA, USA).

### Western Blotting

Cells were washed with phosphate-buffered saline (PBS), detached by trypsinization, and collected by centrifugation. Cell pellets were lysed in RIPA buffer supplemented with 1× protease and phosphatase inhibitor cocktails on ice for 1 h. Lysates were centrifuged at 13,000 rpm for 5 min at 4 °C to remove cellular debris. Protein concentrations were determined by measuring absorbance at 280 nm using a Cytation 5 Cell Imaging Multimode Reader (BioTek Instruments, Winooski, VT, USA) equipped with a microvolume measurement plate. Briefly, 2 µL of each lysate was analyzed using RIPA buffer containing the same protease and phosphatase inhibitors as a blank. Protein concentrations were used to normalize sample loading. For each sample, 100 µg of total protein was prepared in RIPA buffer and Laemmli sample buffer (Bio-Rad Laboratories, Hercules, CA, USA) supplemented with β-mercaptoethanol and denatured at 95 °C for 5 min. Subsequently, 20 µL of each sample, corresponding to 20 µg of protein, was loaded onto 10% SDS–PAGE gels (Cat. No. 4568033; Bio-Rad) and separated by electrophoresis using Tris–glycine running buffer. Proteins were transferred onto nitrocellulose membranes using the Trans-Blot Turbo Transfer System (Bio-Rad; 25 V, 2.5 A), and transfer efficiency was verified by Ponceau S staining.

Membranes were blocked for 1 h at room temperature in 5% bovine serum albumin (BSA) prepared in Tris-buffered saline containing Tween-20 (TBST; 25 mM Tris, pH 8.0, 2.7 mM KCl, 137 mM NaCl, and 0.1% Tween-20). Membranes were incubated overnight at 4 °C with primary antibodies diluted 1:1,000 in blocking buffer, except for anti-α-tubulin (1:4,000). Primary antibodies included anti-AMPKα2 (Cat. Nos. 2757S/2757P), anti-hexokinase II (HK2; Cat. No. 2867S), anti-hexokinase I (HK1; Cat. No. 2024T), anti-AMPKα1 (Cat. No. 2795S), anti-LDHA (clone C4B5), anti-PKM1/2 (clone C103A3), anti-PFKP (clone DAB2), anti-pyruvate dehydrogenase (clone C54G1), anti-S6 (Cat. No. 2317S), and anti-phospho-S6 (Cat. No. 4858S), all from Cell Signaling Technology (Danvers, MA, USA); anti-GFP (Cat. No. ab6556), anti-GLUT1 (Cat. No. ab652), anti-TRIM28/KAP1 (Cat. No. ab22553), and anti-MAGEA3 (clone EPR19065; Cat. No. ab223162) from Abcam (Cambridge, UK); anti-GFP (Cat. No. A11122; Invitrogen); and anti-α-tubulin (Cat. No. T5168) and anti-β-actin (Cat. No. A5441) from Sigma-Aldrich (St. Louis, MO, USA). Following three washes with TBST, membranes were incubated for 1 h at room temperature with the appropriate horseradish peroxidase (HRP)-conjugated secondary antibodies, goat anti-rabbit IgG–HRP (Cat. No. 7074P2) or horse anti-mouse IgG–HRP (Cat. No. 7076S), both from Cell Signaling Technology, diluted 1:10,000. After additional washes with TBST, protein bands were detected using Amersham ECL Western Blotting Detection Reagents (Cat. No. RPN2209; Cytiva, Buckinghamshire, UK) and visualized using an Azure 600 Imaging System (Azure Biosystems, Dublin, CA, USA; software version 1.6.2.1020). Band intensities were quantified by densitometric analysis using Fiji/ImageJ and normalized to α-tubulin, β-actin, or TRIM28, as appropriate. Western blot experiments were performed in at least two independent biological replicates unless otherwise indicated.

### Colony Formation Assay

Following siRNA transfection, cells were harvested, counted, and seeded in duplicate at densities of 500 or 1,000 cells per well in 6-well plates. Cells were cultured for 14 days to allow colony formation, with complete growth medium replaced every 2–3 days. Colonies were fixed and stained for 20 min using crystal violet solution (0.05% w/v crystal violet, 1% formaldehyde, and 1% methanol in PBS), washed with water, air-dried, and imaged using an Azure 600 Imaging System (Azure Biosystems, Dublin, CA, USA). Colony formation was quantified using Fiji/ImageJ (version 1.54k) with the ColonyArea plugin [59]. Images were rotated, converted to 8-bit grayscale, and individual wells were extracted for analysis. Manual thresholding was applied to distinguish stained colonies from background, followed by conversion to binary images. Colonies were identified using the Analyze Particles function with a size threshold of 0–Infinity pixels² and a circularity range of 0.00–1.00. Detected colonies were visually verified using particle outlines and quantified for each well. Colony formation was expressed as a percentage relative to the siRev-L1 control, and data were analyzed and visualized using GraphPad Prism (GraphPad Software, Boston, MA, USA).

### Wound Healing Assay

Following siRNA transfection, DAN-G cells (2.5 × 10⁵ cells per well) were seeded in 24-well plates with complete growth medium. After 24 h, a uniform scratch was created across the cell monolayer in each well using a sterile pipette tip. Detached cells and debris were gently removed, and fresh medium was added. Plates were returned to the incubator, and wound closure was monitored at 24-h intervals. Representative images were captured using an EVOS inverted microscope to assess cell migration. Wound-healing images were analyzed using Fiji/ImageJ. Images were converted to 8-bit grayscale and processed using a fast Fourier transform (FFT) bandpass filter with the autoscale and saturate image options enabled. Image thresholding was subsequently adjusted to distinguish the wound area from the surrounding cell-covered region, followed by application of a minimum filter with a radius of 7 pixels, as previously described for manual wound-area analysis using ImageJ/Fiji [60]. The wound region was delineated using the selection tool, and wound area was measured at 0 and 24 h. Percentage wound closure was calculated relative to the initial wound area using the formula: [(wound area at 0 h − wound area at 24 h)/wound area at 0 h] × 100, consistent with established scratch-assay quantification approaches [61]. Statistical analyses and final data visualization were performed using GraphPad Prism. Data distributions were assessed using the D’Agostino–Pearson, Anderson–Darling, Shapiro–Wilk, and Kolmogorov–Smirnov normality tests. Differences in percentage wound closure between the two experimental groups were analyzed using an unpaired t-test with Welch’s correction, which does not assume equal variances between groups.

### Migration assay

DAN-G cells were treated with siRNAs as described previously. Following treatment, 2.5 × 10⁵ cells were seeded into the apical chamber of Transwell inserts in 500 µL of serum-free medium, while 1.5 mL of complete medium containing FBS was added to the basal chamber as a chemoattractant. Cells were incubated for 24 h to allow migration. Following incubation, non-migrated cells remaining on the upper surface of the membrane were gently removed with a cotton swab. Cells that had migrated to the lower surface of the membrane were fixed, stained, and imaged by automated microscope (ECHO, San Diego, CA, USA) [62].

## RESULTS

### MAGEA3/6 Selectively Promotes Resistance to 2-DG-Induced Glycolytic Stress in Pancreatic Cancer Cells

Our previous study demonstrated, for the first time, that *MAGEA* genes, predominantly expressed in the male germline and frequently reactivated in human cancers as cancer-testis antigens (CTAs), protect spermatogenesis during prolonged food deprivation. Consistently, spermatogonial stem cells from *Magea* knockout mice exhibited increased sensitivity to glycolytic inhibition by 2-deoxyglucose (2-DG). This protective function extended to cancer cells, where MAGEA6 expression enhanced the sustained growth of MIA PaCa-2 pancreatic cancer cells under 2-DG-induced glycolytic stress [14]. The present study investigates the molecular mechanisms underlying MAGEA3/6-mediated resistance to 2-DG and its role in metabolic adaptation in pancreatic cancer cells. To address this question, we applied complementary gain- and loss-of-function models, pharmacological inhibitors targeting multiple metabolic pathways, and a combination of cancer cell growth, viability, and metabolic assays, integrated with transcriptomic and multi-omics analyses to elucidate the molecular mechanisms underlying MAGEA3/A6-mediated metabolic adaptation and resistance to 2-DG.

Figure 1a presents a schematic overview of glucose metabolism, highlighting glycolysis, the tricarboxylic acid (TCA) cycle, and interconnected metabolic pathways, including the pentose phosphate pathway (PPP), hexosamine biosynthetic pathway (HBP), and glutamine metabolism feeding into the TCA cycle, along with the specific metabolic inhibitors used in this study. We targeted glycolysis using 2-deoxyglucose (2-DG), glycolysis and mitochondrial metabolism using 3-bromopyruvate (3-BP), the pentose phosphate pathway (PPP) using 6-aminonicotinamide (6-AN), and glutamine metabolism using BPTES, a selective inhibitor of glutaminase 1 (GLS1) [63]. MIA PaCa-2 cells expressing MAGEA3/6 were treated with increasing concentrations of 2-DG and exhibited a rightward shift in the dose–response curve compared with GFP-expressing control cells, resulting in a more than 2.5-fold increase in the half-maximal inhibitory concentration (IC50). These findings demonstrate that MAGEA3/6 expression enhances resistance to 2-DG-induced metabolic stress, consistent with our previously published findings (Fig. 1b) [64]; thus growth characteristics were determined at different seeding densities. GFP and A3/6 cells exhibited similar basal growth patterns over 3 days suggesting that differences in basal cell growth were not likely to account for the observed difference in 2-DG sensitivity (Fig. 1c) alone. To confirm this phenotype, additional gain-of-function models of MIA PaCa-2 expressing empty vector, MAGEA3, or MAGEA6 were generated. MAGE-expressing cells were more resistant to 2-DG compared to empty-vector controls. Western blot confirmed expression of MAGEA3 and MAGEA6 (Fig. 1d–e). MAGEA6 was also examined in PANC-1-Luc2 cells, where MAGEA6 expression also changed 2-DG sensitivity compared with empty-vector controls (Fig.1f). In this model, RT-qPCR validated the increased expression of MAGEA6 (Fig. 1g). Additionally, we investigated the effect of MAGEA3/6 expression on the response of HEK293 cells to 2-DG to determine whether the MAGE-associated resistance phenotype was specific to cancer cells or could also be observed in a non-cancerous cellular background. Western blotting confirmed the expression of MAGEA3 and MAGEA6 in the HEK293 cells. We didn’t see a significant difference between MAGEA3 and A6 cells compared to empty vector (Fig. S1a-b). Together, these experiments demonstrated a reproducible association between MAGEA3/6 expression and cellular tolerance to glycolytic inhibition by 2-DG in independently generated models.

**Figure 1.**
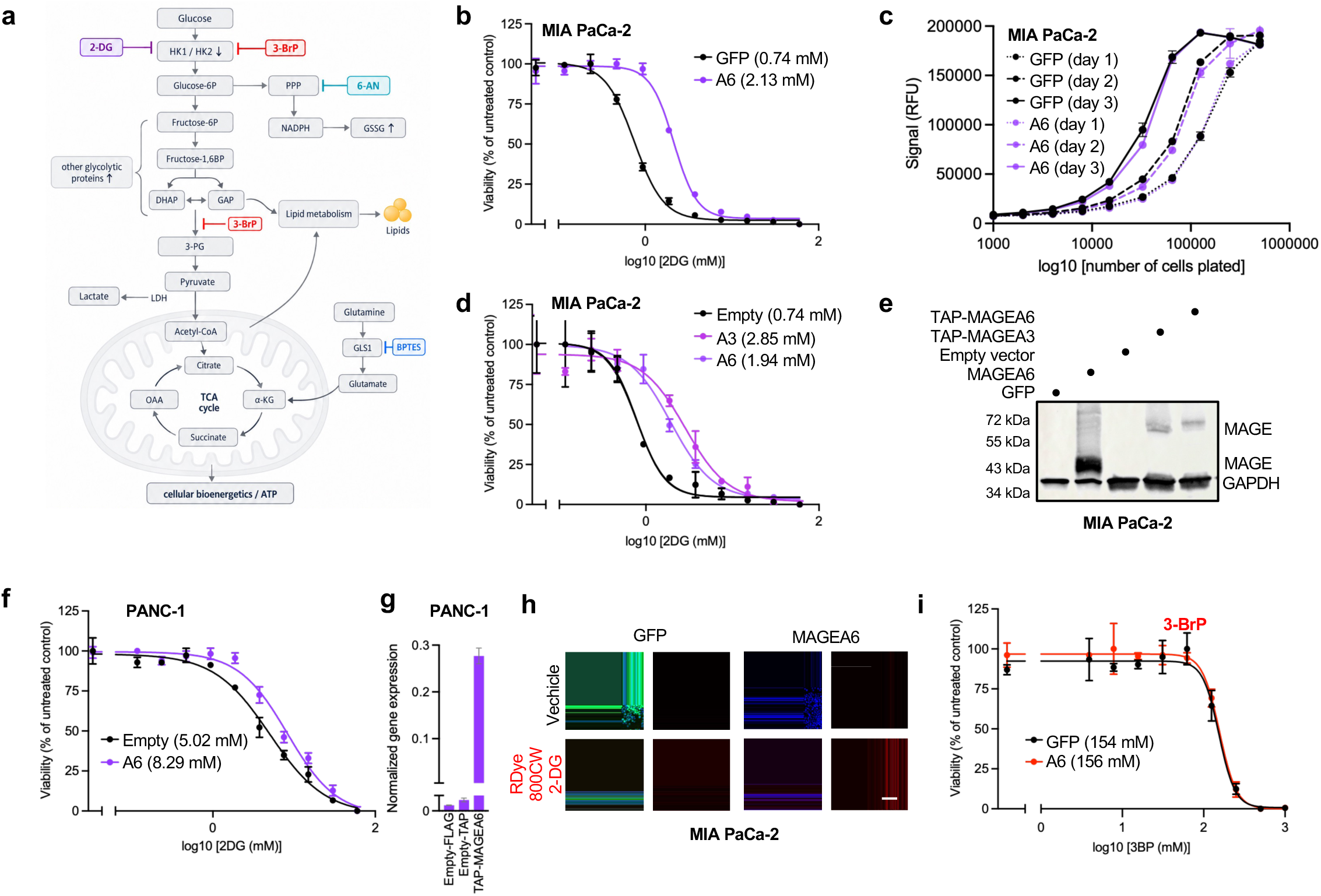
MAGEA6 promotes resistance to 2-DG-induced metabolic stress in pancreatic cancer cells. **a.** Schematic representation of glucose metabolism, highlighting glycolysis, the pentose phosphate pathway, mitochondrial metabolism, and selected sites of pharmacologic inhibition, including 2-deoxy-D-glucose (2-DG), 3-bromopyruvate (3-BrP), and 6-aminonicotinamide (6-AN). **b.** Dose-response curves showing the sensitivity of GFP-control and MAGEA6 (A6)-expressing MIA PaCa-2 cells to 2-DG. Cell viability was measured using the Alamar Blue (AB) assay 3 days after treatment. **c.** Growth curves of GFP-control and MAGEA6-expressing MIA PaCa-2 cells plated at increasing cell numbers per well in 96-well plates and analyzed using the AB assay on days 1–3, demonstrating the dynamic range of the assay and comparable baseline growth between the two cell lines. **d.** Dose-response curves showing the sensitivity of newly generated MIA PaCa-2 gain-of-function (GOF) models expressing empty vector, MAGEA3 (A3), or MAGEA6 (A6) to 2-DG, measured using the AB assay 5 days after treatment. **e.** Western blot analysis confirming MAGEA3 and MAGEA6 expression in the MIA PaCa-2 GOF models. **f.** Dose-response curves showing the sensitivity of PANC-1 cells expressing empty vector or MAGEA6 to 2-DG. **g.** RT-qPCR analysis of MAGEA6 expression in PANC-1 cells, normalized to 18S rRNA. **l.** Representative fluorescence images of GFP-control and MAGEA6-expressing MIA PaCa-2 cells treated with vehicle or IRDye 800CW 2-DG (5 μM), illustrating cellular 2-DG uptake. **m.** Dose-response curves showing the sensitivity of GFP-control and MAGEA6-expressing MIA PaCa-2 cells to the glycolytic inhibitor 3-BrP, measured 5 days after treatment. All dose-response curves assessing sensitivity to 2-DG or 3-BrP were generated using the Alamar Blue assay. Data are presented as viability (% of untreated control), with IC₅₀ values indicated in mM for the corresponding compound. Data represent mean ± SD of four technical replicates; experiments were repeated at least three times, and one representative experiment is shown.

Next, we asked whether MAGEA6-mediated resistance is modulated by the availability of extracellular glucose. To evaluate the fitness effects of MAGEA6 expression in the setting of glucose-associated metabolic stress, we performed longitudinal analysis across increasing concentrations of 2-DG and decreasing concentrations of glucose, MAGEA6-expressing cells maintained greater resistance to higher concentrations of 2-DG over time, whereas no substantial differences in growth patterns were observed between MAGEA6-expressing and GFP-control cells across different glucose concentrations (Fig. S1c). Intriguingly, even under glucose-free conditions, MAGEA6-expressing cells exhibited approximately 1.6-fold (60%) greater resistance to 2-DG treatment than GFP-expressing controls, although this protective effect was less pronounced than under glucose-replete conditions (Fig. S1d). These findings suggest that MAGEA6-mediated resistance is not exclusively dependent on glucose availability but may also involve alternative metabolic adaptations or glycolysis-independent cytoprotective mechanisms. We next investigated whether MAGEA3/6 expression influences cellular 2-DG uptake by assessing the uptake of the fluorescent analog IRDye 800CW 2-DG in MIA PaCa-2 cells expressing either GFP or MAGEA3/6 (Fig.1h). Finally, to determine whether MAGEA3/6-mediated resistance extends beyond glycolytic inhibition by 2-DG, we treated cells with 3-bromopyruvate (3-BrP), a multifunctional metabolic inhibitor that disrupts glycolysis through inhibition of HK2 while also impairing mitochondrial respiration and ATP production (Fig. 1a). Unlike 2-DG, 3-bromopyruvate (3-BrP) treatment did not reveal a significant difference in sensitivity between MAGEA3/6-expressing and GFP-control cells (Fig. 1i). Together, these findings suggest that MAGEA3/6 selectively enhances pancreatic cancer cell tolerance to 2-DG-induced metabolic stress rather than conferring broad resistance to glycolytic inhibition or nutrient deprivation [65].

### MAGEA3/6 Depletion Reduces Pancreatic Cancer Cell Fitness and Sensitizes Cells to 2-DG-Induced Metabolic Stress

We next wanted to complement our gain-of-function studies by analyzing whether endogenous expression of MAGEA3/6 is required for pancreatic cancer cell growth and adaptation to metabolic stress. Analysis of the Cancer Cell Line Encyclopedia (CCLE) data revealed significant heterogeneity of MAGE family expression in pancreatic cancer cell lines. BXPC-3 and DAN-G were among the cells with high expression of MAGEA3/6 and picked as loss of function models (Fig. 2a). Differential endogenous MAGEA3/6 expression was confirmed by RT-qPCR analysis of selected cell lines and used to identify MAGEA3/6 positive models for further loss-of-function experiments (Fig. 2b). MAGEA3/6 expression was then decreased by siRNA. The knockdown efficiency was verified by RT-qPCR in selected pancreatic cancer cell lines (Fig. 2c) and by Western blotting in BxPC-3 cells (Fig. 2d). Next, we investigated the impact of endogenous MAGEA3/6 depletion on the long-term proliferative potential of pancreatic cancer cells. Knockdown of MAGEA3/6 reduced colony formation in BxPC-3 cells compared with siRNA control siRevL1, which suggests impaired clonogenic growth after loss of MAGEA3/6 (Fig. 2e–f). To see whether this phenotype could be recapitulated in an independent model of MAGEA3/6-positive pancreatic cancer, loss-of-function experiments were performed in DAN-G cells. Western blot showed decreased expression of MAGEA3/6 after siRNA treatment (Fig. 2g). MAGEA3/6 depletion significantly inhibited colony formation compared to control cells (Fig. 2h–i). In all, these findings support a role for endogenous MAGEA3/6 in promoting the proliferative and clonogenic capacity of pancreatic cancer cells.

**Figure 2.**
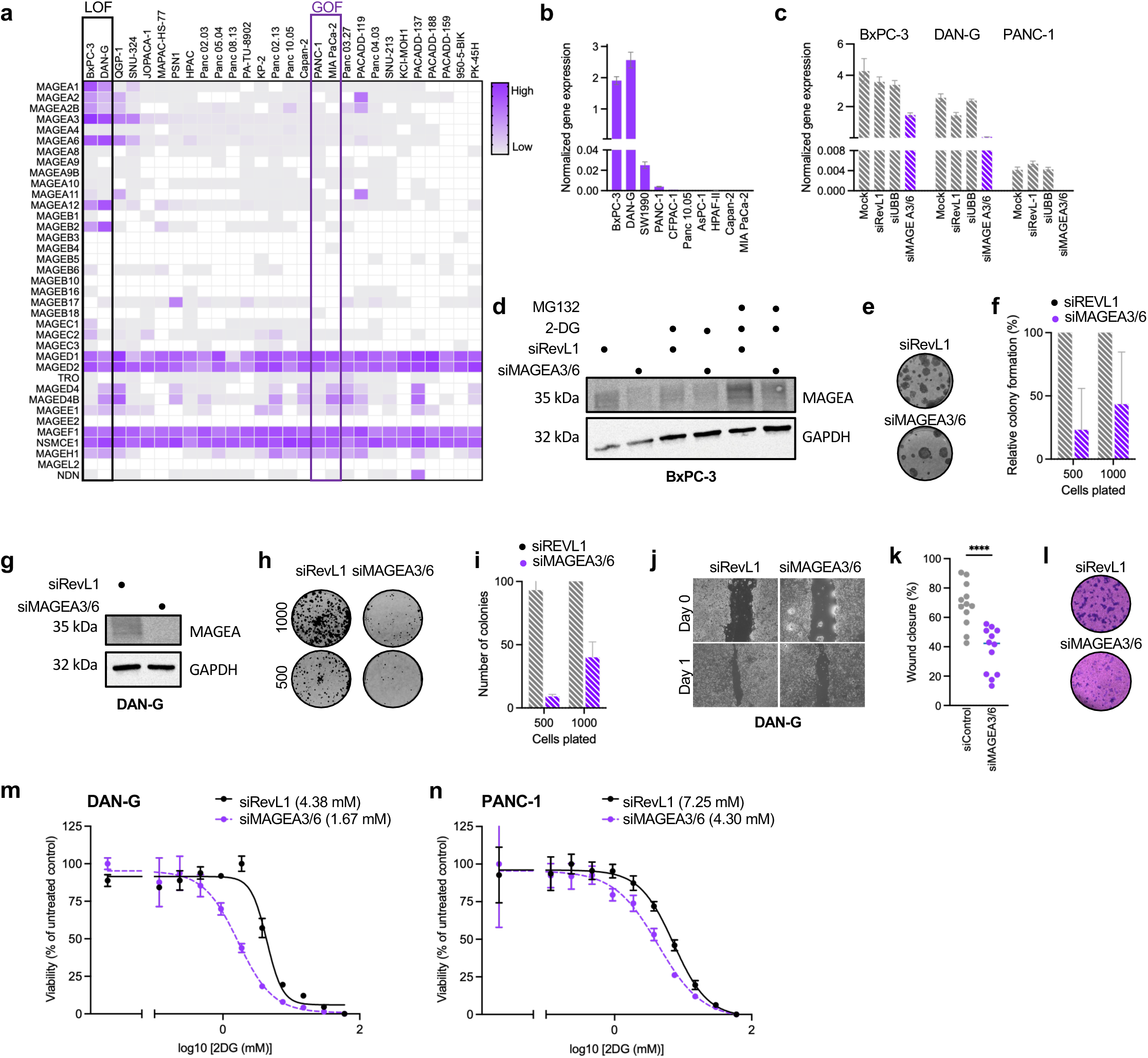
MAGEA6 is required for pancreatic cancer cell survival and resistance to metabolic stress. **a.** Heatmap showing MAGE gene expression across pancreatic cancer cell lines; data were obtained from the Cancer Cell Line Encyclopedia (CCLE) on August 24, 2022. **b.** RT-qPCR analysis of MAGEA3/A6 expression in selected pancreatic cancer cell lines. **c.** RT-qPCR analysis of MAGEA3/A6 expression following siRNA-mediated knockdown in MAGEA3/A6-positive pancreatic cancer cells. **d.** Western blot analysis of MAGEA3/A6 expression following siRNA-mediated knockdown in MAGEA3/A6-positive BxPC3 cells. **e.** Representative colony-formation assays and **f.** corresponding quantification showing the effect of MAGEA3/A6 knockdown on clonogenic growth in BxPC3 cells under the indicated conditions. **g.** Western blot analysis of MAGEA3/A6 expression following siRNA-mediated knockdown in DAN-G cells. **h.** Representative colony-formation assay image and **i.** corresponding quantification following MAGEA3/A6 knockdown in DAN-G cells. **j.** Representative images of scratch-wound assays and **k.** quantification of wound closure following MAGEA3/A6 knockdown in DAN-G cells. Statistical significance for wound closure was determined using Welch’s *t*-test. **l.** Representative microscopy images of migration assays in DAN-G cells following MAGEA3/A6 knockdown. **m.** Dose-response curves showing the sensitivity of control and MAGEA3/A6-knockdown DAN-G cells to 2-deoxy-D-glucose (2-DG), measured using the Alamar Blue assay 5 days after treatment. **n.** Dose-response curves showing the sensitivity of control and MAGEA3/A6-knockdown PANC-1 cells to 2-DG, measured using the Alamar Blue assay 5 days after treatment. Data are presented as viability (% of untreated control), with IC₅₀ values indicated in mM 2-DG. Data represent mean ± SD of four technical replicates; experiments were repeated at least three times, and one representative experiment is shown. Statistical significance was determined using the indicated statistical tests; *P* < 0.05, **P** < 0.01, ***P*** < 0.001, and **P** < 0.0001.

Since tumor progress is also affected by cancer cell motility [66], we next determined the effect of MAGEA3/6 depletion on DAN-G cell migration. Scratch-wound analysis showed that knockdown of MAGEA3/6 reduced wound closure compared with control cells (Fig. 2j–k). Consistent with this observation, migration assays showed less migrated cells upon MAGEA3/6 depletion (Fig. 2l). These findings imply a role for endogenous MAGEA3/6 in clonogenic growth and the migratory phenotype of pancreatic cancer cells. Eventually, we investigated the role of endogenous MAGEA3/6 in the resistance to glycolytic stress. Control or MAGEA3/6-knockdown DAN-G cells were treated with increasing concentrations of 2-DG. MAGEA3/6 depletion enhanced the sensitivity to 2-DG, as shown by the dose-response profiles and the corresponding IC50 values (Fig. 2m). A similar experiment was carried out in PANC-1 cells and MAGEA3/6 knockdown also changed the response to 2-DG relative to control cells (Fig. 2n). Together, these complementary loss- and gain-of-function findings support a role for MAGEA3/6 in maintaining pancreatic cancer cell fitness and promoting resistance to 2-DG-induced glycolytic stress.

### MAGEA6 mediates 2-DG resistance independent of AMPK-mTOR signaling

Given the increased dependence of many cancer cells on aerobic glycolysis (the Warburg effect), 2-DG initially emerged as a promising anticancer therapeutic [67, 68]. However, its limited clinical efficacy highlighted the remarkable ability of cancer cells to adapt to glycolytic inhibition, even in highly glucose-dependent cancers such as PDAC. This adaptive resistance was subsequently linked to mTORC1-dependent metabolic reprogramming, which enables cancer cells to bypass glycolytic blockade and sustain energy production and growth. Importantly, combined inhibition of mTOR signaling and glycolysis suppressed cancer cell growth, demonstrating that targeting compensatory metabolic pathways can overcome resistance to glycolytic inhibition [48]. This was particularly interesting and attracted our attention, as previous studies have demonstrated that MAGEA3/6 interact with the E3 ubiquitin ligase TRIM28 to promote the ubiquitination and degradation of AMPKα1, a master regulator of cellular energy homeostasis and metabolic adaptation during nutrient stress. AMPK and mTORC1 function as opposing regulators of cellular metabolism, coordinating catabolic and anabolic processes in response to nutrient and energy availability. Through AMPKα1 degradation, MAGEA3/6 was shown to modulate AMPK–mTOR signaling and cellular responses to metabolic stress [33, 39]. Based on these findings, we hypothesized that MAGEA3/6-mediated resistance to 2-DG-induced glycolytic stress might be driven by suppression of AMPK activity and consequent modulation of mTORC1 signaling. Previous studies by our group and others demonstrated that MAGEA3/6 promote AMPKα1 ubiquitination and degradation and suppress AMPK signaling across multiple cancer types, including lung, breast, colorectal, renal, and brain cancers [33, 38, 69–71]. Given our observation that MAGEA3/6 expression enhances resistance to 2-DG-induced metabolic stress in PDAC cells, we next investigated whether this protective phenotype involves modulation of the AMPK–mTOR signaling axis. We first examined AMPKα1 protein levels in MAGEA6-expressing and GFP-control MIA PaCa-2 cells following long-term adaptation to glycolytic inhibition. Cells were cultured for five weeks in medium containing 2 mM 2-DG, a concentration approximating their IC50, which resulted in increased resistance to subsequent 2-DG treatment (Fig. 1b and S1c). To distinguish the effects of long-term adaptation from acute metabolic stress, both 2-DG-adapted and naïve cells were subsequently challenged with 3 mM 2-DG for 1–24 h, and components of the AMPK–mTOR signaling pathway were examined by Western blotting (Fig. 3a–d).

**Figure 3.**
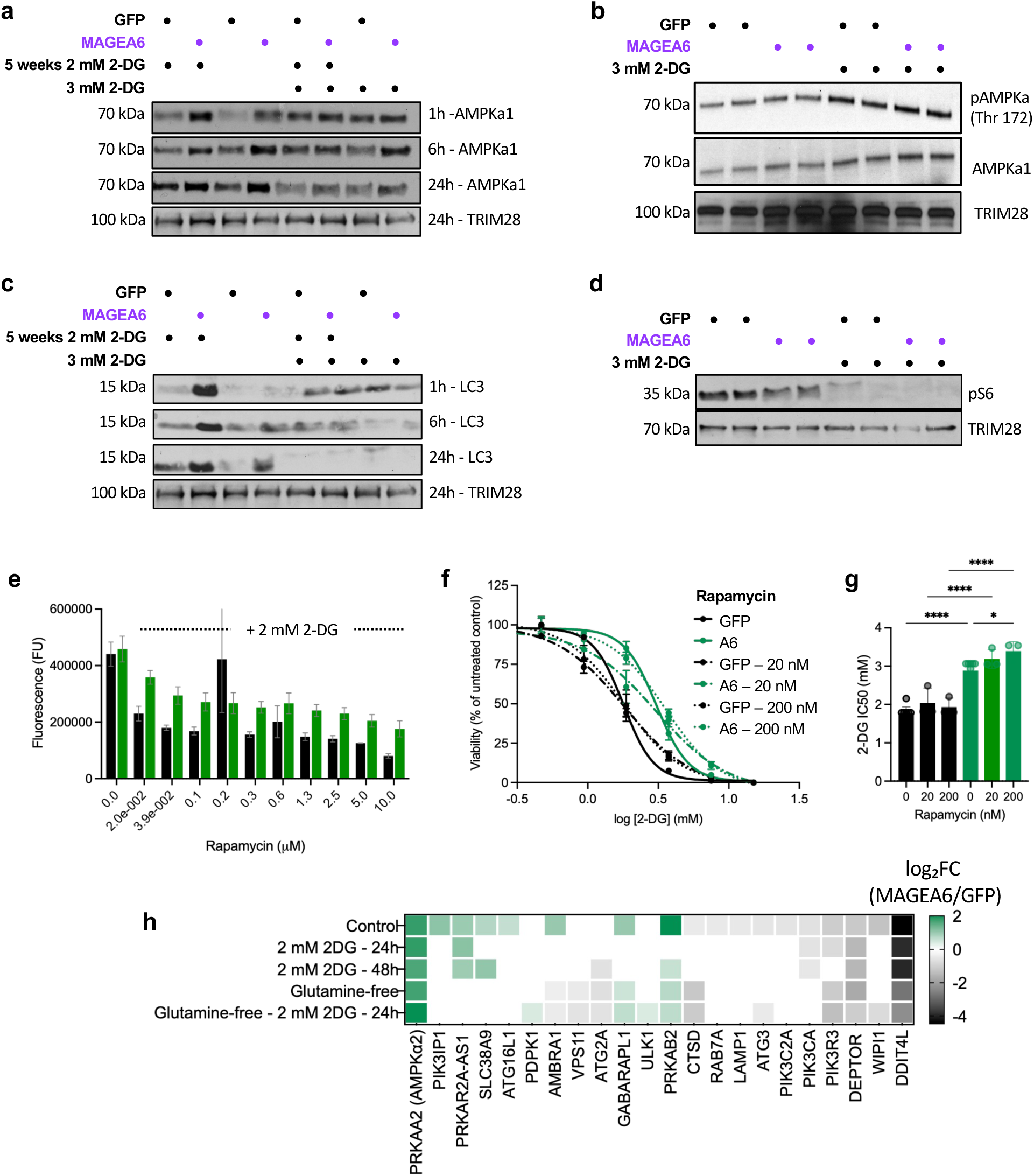
MAGEA6-mediated resistance to 2-DG is largely independent of AMPK–mTOR signaling in MIA PaCa-2 pancreatic cancer cells. a–d. Western blot analysis of components of the AMPK–mTOR signaling pathway in GFP-control and MAGEA6-expressing MIA PaCa-2 cells under basal conditions, following long-term adaptation to 2 mM 2-DG for 5 weeks, or after acute treatment with 3 mM 2-DG for 1–24 h, as indicated. Representative immunoblots show **a.** AMPKα1; **b.** phosphorylated AMPKα (Thr172), total AMPKα, and TRIM28; **c.** LC3; and **d.** phosphorylated S6 (p-S6), together with the corresponding loading controls. **e.** Cell viability of GFP-control and MAGEA6-expressing MIA PaCa-2 cells following 5 days of treatment with increasing concentrations of rapamycin in combination with 2 mM 2-DG, assessed using the Alamar Blue assay and expressed as fluorescence units. **f.** Representative 2-DG dose–response curves for GFP-control and MAGEA6-expressing MIA PaCa-2 cells in the presence of 0, 20, or 200 nM rapamycin. **g.** Quantification of 2-DG half-maximal inhibitory concentration (IC50) values from three independent experiments in GFP-control and MAGEA6-expressing MIA PaCa-2 cells treated with 0, 20, or 200 nM rapamycin. Data are presented as mean ± SD. Statistical significance was assessed by one-way ANOVA followed by Šidák’s multiple-comparisons test, as indicated. \*\*\**P* < 0.0001. **h.** Heatmaps showing log₂ fold changes (MAGEA6/GFP) in the expression of selected genes associated with AMPK–mTOR signaling and autophagy, as determined by RNA-seq analysis.

Unexpectedly, in contrast to previous reports of MAGEA3/6-mediated AMPKα1 degradation, we observed higher AMPKα1 protein abundance in MAGEA6-expressing cells than in GFP controls. Following long-term adaptation to 2-DG, AMPKα1 levels were elevated in MAGEA6-expressing cells compared with GFP controls (Fig. 3a and 3b). In naïve cells, MAGEA6 expression was also associated with modestly increased AMPKα1 abundance, with differences becoming more pronounced following acute 2-DG treatment. Specifically, AMPKα1 protein levels were higher in MAGEA6-expressing cells at 1 and 24 h after treatment (Fig. 3a). To further investigate AMPK signaling, we examined AMPKα phosphorylation at Thr172, a key regulatory site associated with AMPK activation, together with total AMPKα and TRIM28 protein expression (Fig. 3b). These data showed that changes in AMPKα1 abundance were accompanied by alterations in its activation state.

We next examined microtubule-associated protein 1 light chain 3 (LC3), an autophagy-associated protein, because AMPK activation can promote autophagy during metabolic stress [72]. In line with AMPK levels, LC3 protein abundance was markedly elevated in MAGEA6-expressing cells following long-term adaptation to 2-DG across all three examined time points, whereas no substantial differences were observed between MAGEA6 and GFP-control cells following acute treatment without prior adaptation (Fig. 3c). Moreover, LC3 abundance decreased following acute 2-DG treatment in adapted cells, suggesting alterations in autophagy-associated processes. However, because LC3-I and LC3-II could not be reliably resolved, these changes could not be directly attributed to altered autophagic flux. To evaluate downstream mTORC1 signaling, we also examined phosphorylation of ribosomal protein S6, a downstream marker of mTORC1 pathway activity, under the corresponding treatment conditions (Fig. 3d). Together, these observations suggest that MAGEA6-mediated regulation of AMPK signaling is context-dependent and potentially tissue-specific, prompting us to investigate whether pharmacological inhibition of mTORC1 could overcome the protective phenotype conferred by MAGEA6.

To functionally assess the contribution of mTOR signaling, we treated GFP-control and MAGEA6-expressing cells with the mTORC1 inhibitor rapamycin. Rapamycin alone had no appreciable effect on cell viability (Fig. S2a and S2b). However, in combination with 2 mM 2-DG, rapamycin reduced cell viability in both cell lines, with a more pronounced effect in GFP-control cells than in MAGEA6-expressing cells (Fig. 3e). Consequently, MAGEA6-expressing cells maintained a relative survival advantage over GFP controls despite pharmacological inhibition of mTORC1 (Fig. S2c), suggesting that mTORC1 inhibition was insufficient to abolish MAGEA6-mediated resistance to 2-DG-induced metabolic stress. We next more directly examined the effect of mTORC1 inhibition on 2-DG sensitivity by generating complete 2-DG dose response curves in the presence of 0, 20 or 200 nM rapamycin, MAGEA6 cells were more viable compared to GFP control (Fig. 3f). In addition, we calculated and visualized IC50 and confirmed that MAGEA6-expressing cells kept higher 2-DG IC50s than GFP-control cells even in the presence of rapamycin (Fig. 3g). We also compared the fold change in IC50 upon treatment with 20 or 200 nM rapamycin compared to 2-DG treatment alone withing each genotype, which showed that the differential sensitivity to 2-DG between the two cell populations was not abolished by pharmacological inhibition of mTORC1 (Fig. S2d).

To get better insight into the molecular mechanisms underlying MAGEA6-mediated resistance to 2-DG, we performed integrated RNA-seq, proteomic, metabolomic, and lipidomic analyses of GFP-control and MAGEA6-expressing MIA PaCa-2 cells cultured in complete or glutamine-free medium and treated with 2 mM 2-DG for 24 or 48 h, as indicated (Fig. S3a). Three biological replicates were analyzed per condition. Hierarchical clustering (Fig. S3b) and principal component analysis (PCA; Fig. S3c) of RNA-seq data revealed distinct transcriptional profiles driven primarily by MAGEA6 expression and glutamine availability, whereas the effects of 2-DG treatment were less pronounced. These findings suggest that MAGEA6 influences transcriptional adaptations to nutrient availability and metabolic stress. We first analyzed genes involved in AMPK–mTOR signaling (Fig. 3h). Transcriptomic analysis revealed consistent upregulation of *PRKAA2* (AMPKα2) and downregulation of the mTOR regulators *DDIT4L* and *DEPTOR* in MAGEA6-expressing cells across all metabolic conditions. Together with our Western blot analyses, these findings suggest that MAGEA6-mediated regulation of AMPK–mTOR signaling is context-dependent and that resistance to 2-DG does not require AMPK downregulation. Furthermore, pharmacological inhibition of mTORC1 failed to abolish the MAGEA6-associated resistance advantage, indicating that the protective phenotype cannot be explained solely by the canonical MAGEA3/6–AMPK–mTOR axis and likely involves additional metabolic adaptations that support pancreatic cancer cell survival under glycolytic stress.

### Pentose phosphate pathway inhibition enhances sensitivity to 2-DG independently of genotype, while MAGEA6-expressing cells retain greater resistance

Similar to glucose, 2-DG is phosphorylated by hexokinase to 2-DG-6-phosphate, which accumulates intracellularly and inhibits glycolysis, in part through competitive inhibition of glucose-6-phosphate isomerase (GPI), leading to reduced levels of downstream glycolytic intermediates, including fructose-1,6-bisphosphate. Interestingly, glycolysis-independent cancer cells exhibit relatively higher levels of PPP metabolites even under untreated conditions, a difference that becomes more pronounced following 2-DG treatment, consistent with increased diversion of glucose-6-phosphate into the PPP [48]. PPP is a major metabolic branch of glucose metabolism that utilizes glucose-6-phosphate to generate NADPH and ribose-5-phosphate, supporting redox homeostasis, glutathione reduction, and nucleotide biosynthesis [73]. Because 2-DG inhibits glycolysis and may promote the diversion of glucose-6-phosphate into the PPP, we next investigated whether MAGEA6-mediated resistance to glycolytic stress involves increased reliance on this alternative metabolic pathway. To test this hypothesis, we used 6-aminonicotinamide (6-AN), an inhibitor of the oxidative PPP, to determine whether PPP inhibition could overcome MAGEA6-associated resistance to 2-DG. A schematic representation of glycolysis and the PPP, highlighting pharmacological inhibition of the PPP by 6-AN, is shown in Fig. 4a.

**Figure 4.**
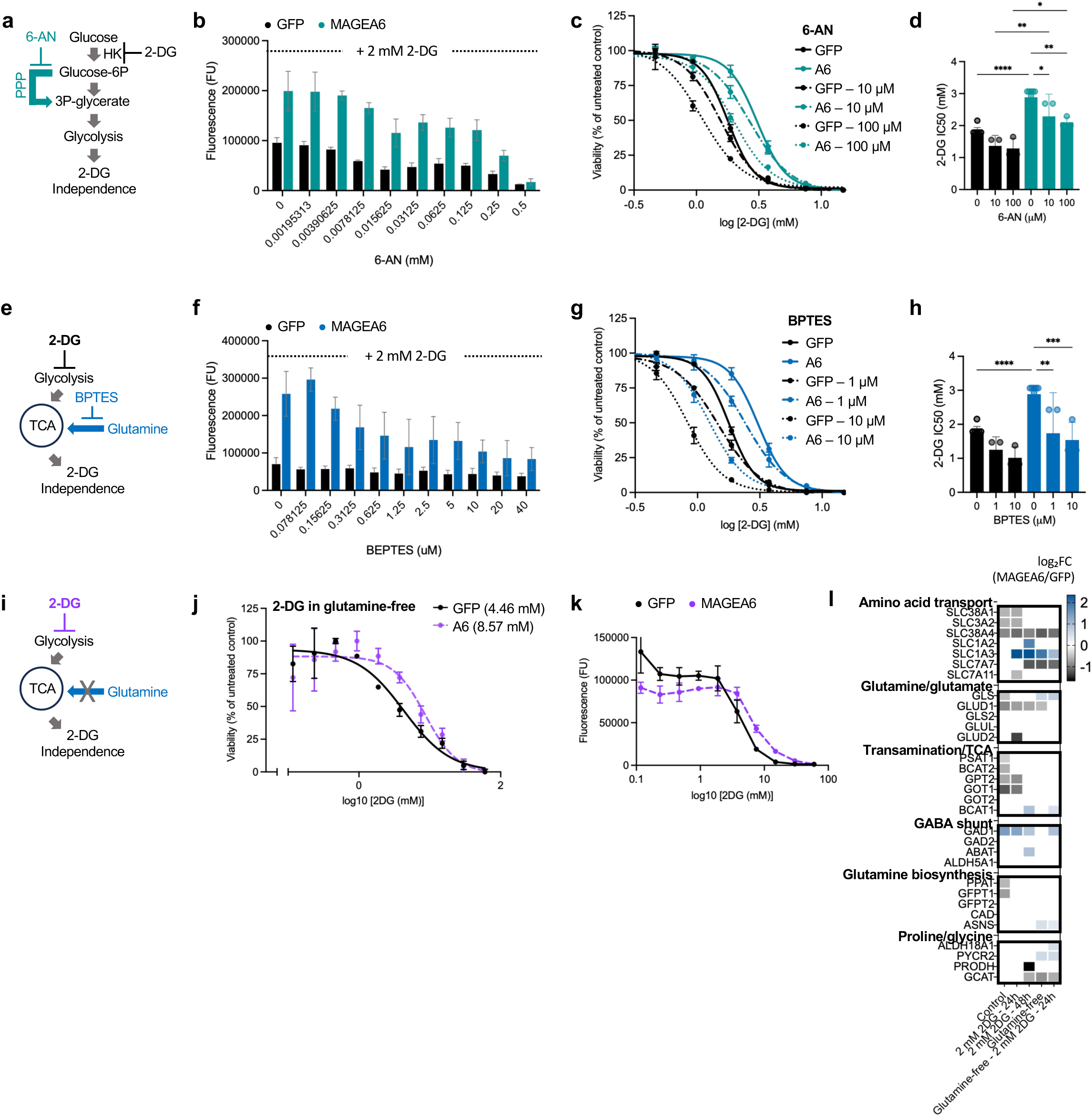
MAGEA6-expressing MIA PaCa-2 pancreatic cancer cells retain resistance to 2-DG despite inhibition of the pentose phosphate pathway, glutaminase activity, and glutamine deprivation. **a.** Schematic representation of glycolysis and the pentose phosphate pathway (PPP), highlighting pharmacological inhibition of the PPP by 6-aminonicotinamide (6-AN). **b.** Viability of GFP-control and MAGEA6-expressing MIA PaCa-2 cells following 5 days of treatment with increasing concentrations of 6-AN in combination with 2 mM 2-DG, assessed using the Alamar Blue assay and expressed as fluorescence units (FU). **c.** Representative 2-DG dose–response curves for GFP-control and MAGEA6-expressing MIA PaCa-2 cells treated with increasing concentrations of 2-DG in the presence of 0, 10, or 100 µM 6-AN. Cell viability was assessed using the Alamar Blue assay after 5 days of treatment. **d.** Quantification of 2-DG half-maximal inhibitory concentration (IC₅₀) values from three independent experiments in GFP-control and MAGEA6-expressing cells treated with the indicated concentrations of 6-AN. **e.** Schematic representation of glutamine metabolism and its pharmacological inhibition by the glutaminase inhibitor BPTES. **f.** Viability of GFP-control and MAGEA6-expressing MIA PaCa-2 cells following 5 days of treatment with increasing concentrations of BPTES in combination with 2 mM 2-DG, assessed using the Alamar Blue assay and expressed as fluorescence units (FU). **g.** Representative 2-DG dose–response curves for GFP-control and MAGEA6-expressing cells treated with increasing concentrations of 2-DG in the presence of 0, 1, or 10 µM BPTES. Cell viability was assessed using the Alamar Blue assay after 5 days of treatment. **h.** Quantification of 2-DG IC50 values from three independent experiments in GFP-control and MAGEA6-expressing cells treated with the indicated concentrations of BPTES. **i.** Schematic representation of glutamine deprivation as an alternative approach to restricting glutamine metabolism. **j.** 2-DG dose–response curves for GFP-control and MAGEA6-expressing MIA PaCa-2 cells cultured in glutamine-free medium, showing cell viability normalized to the corresponding untreated controls and used to determine 2-DG IC₅₀ values. **k.** Corresponding 2-DG dose–response curves showing cell viability in glutamine-free medium, measured using the Alamar Blue assay and expressed as fluorescence units (FU). **l.** RNA-seq heatmap showing differential expression of genes involved in glutamine, glutamate, and interconnected amino acid metabolism in MAGEA6-expressing cells relative to GFP controls under basal conditions, following treatment with 2 mM 2-DG for 24 or 48 h, and under glutamine deprivation with or without 2 mM 2-DG for 24 h. Genes are organized into six functional groups. Heatmap displays log₂ fold changes (MAGEA6/GFP) for genes meeting the differential-expression criteria (adjusted *P* ≤ 0.05 and |log₂FC| ≥ 1) in at least one experimental condition. Within each functional group, genes are ordered by descending log₂ fold change under untreated control conditions, with genes lacking reported control values positioned afterward. Data are presented as mean ± SD, where applicable. Statistical comparisons of IC50 values in panels **d** and **h** were performed using one-way ANOVA followed by Šidák’s multiple-comparisons test. **P < 0.01, ***P < 0.001, ****P < 0.0001.

We first evaluated the dose-dependent effects of 6-AN on GFP-control and MAGEA6-expressing MIA PaCa-2 cells to determine their relative sensitivity and establish an appropriate concentration range for subsequent experiments. Dose–response analysis revealed no appreciable difference in sensitivity to 6-AN between the two cell populations, with both exhibiting comparable reductions in viability across concentrations ranging from 0.001 to 0.5 mM (Fig. S4a). In contrast, combined treatment with 6-AN and 2 mM 2-DG resulted in greater viability of MAGEA6-expressing cells compared with GFP controls (Fig. 4b). This survival advantage was further demonstrated by normalizing the viability of MAGEA6-expressing cells to that of GFP-control cells under the corresponding treatment conditions, revealing consistently approximately twofold higher viability in MAGEA6-expressing cells at 6-AN concentrations up to 0.25 mM (Fig. S4b). These findings indicated that MAGEA6-expressing cells retain their survival advantage even when glycolysis and the PPP are simultaneously challenged. To further investigated whether PPP inhibition changes MAGEA6-mediated resistance to 2-DG, we performed the reciprocal experiment by generating 2-DG dose–response curves in the presence of fixed concentrations of 6-AN (0, 10, or 100 µM; Fig. 4c). Both cells exhibited increased sensitivity to 2-DG in the presence of 6-AN, indicating that simultaneous targeting of glycolysis and the PPP enhances metabolic stress. Nevertheless, MAGEA6-expressing cells maintained greater resistance to 2-DG than GFP controls. Consistent with these observations, quantification of 2-DG IC50 values from three independent experiments confirmed that MAGEA6-expressing cells retained higher IC50 values than GFP-control cells in the presence of 6-AN (Fig. 4d). To determine whether 6-AN differentially affected 2-DG sensitivity between the two cell lines, we compared the fold changes in IC50 values following treatment with 10 or 100 µM 6-AN relative to 2-DG treatment alone within each genotype (Fig. S4c). 6-AN similarly reduced 2-DG IC50 values in both GFP-control and MAGEA6-expressing cells without abolishing the survival advantage conferred by MAGEA6 expression. These findings suggest that MAGEA6-mediated protection against glycolytic stress cannot be explained solely by increased dependence on the PPP and likely involves additional metabolic adaptations. However, the potential off-target effects of 6-AN and the absence of direct PPP flux measurements preclude definitive conclusions about PPP independence.

### MAGEA6-cells maintains resistance to 2-DG despite inhibition of glutamine metabolism

Glycolysis-independent cancer cells can overcome glycolytic inhibition by increasing their reliance on glutamine metabolism, which replenishes TCA cycle intermediates through glutaminolysis, supports mitochondrial energy production, and sustains aspartate biosynthesis and cell proliferation [48]. Therefore, we next investigated whether MAGEA6-mediated resistance to 2-DG depends on glutamine metabolism. To determine whether disruption of this pathway alters sensitivity to glycolytic inhibition, we treated GFP-control and MAGEA6-expressing MIA PaCa-2 cells with BPTES, a selective inhibitor of glutaminase 1 (GLS1), either alone or in combination with 2-DG (Fig. 4e-h). Interestingly, treatment with increasing concentrations of BPTES alone had little effect on the viability of either GFP-control or MAGEA6-expressing cells (Fig. S4d). However, when BPTES was combined with 2 mM 2-DG, cell viability decreased in both populations in a BPTES concentration-dependent manner, with a more pronounced reduction in GFP-control cells (Fig. 4f). Combined treatment reduced viability more substantially than BPTES alone, suggesting that disruption of glutamine metabolism increases cellular vulnerability to glycolytic inhibition. Nevertheless, MAGEA6-expressing cells maintained greater viability than GFP controls across the combined treatment conditions, as further demonstrated by the relative fold change in viability between the two cell populations (Fig. S4e).

To further assess whether glutaminase inhibition alters sensitivity to 2-DG, we generated 2-DG dose–response curves in the presence of 0, 1, or 10 µM BPTES (Fig. 4g). BPTES treatment shifted the dose–response curves toward greater sensitivity to 2-DG in both cell populations, suggesting that glutamine metabolism contributes to cellular adaptation during glycolytic stress. This effect was confirmed by quantification of 2-DG IC50 values from three independent experiments (Fig. 4h). In GFP-control cells, the mean 2-DG IC50 decreased from 1.79 mM without BPTES to 1.47 and 0.83 mM in the presence of 1 and 10 µM BPTES, respectively. Similarly, in MAGEA6-expressing cells, the corresponding IC50 values decreased from 2.92 to 2.43 and 1.24 mM. Thus, both populations became increasingly sensitive to 2-DG as BPTES concentration increased, but MAGEA6-expressing cells consistently maintained approximately 1.5–1.6-fold higher IC50 values than GFP controls, showing that their relative resistance to glycolytic inhibition persisted despite glutaminase inhibition (Fig. 4h). To compare the effects of BPTES on 2-DG sensitivity within each cell population, we normalized the IC50 values obtained under BPTES treatment to those of the corresponding controls without BPTES (Fig. S4f), showing that glutaminase inhibition sensitizes both populations to glycolytic stress. However, the persistent difference in IC50 values between the two populations suggests that MAGEA6-mediated resistance is not abolished by pharmacological inhibition of glutamine metabolism.

To independently investigate the contribution of glutamine availability to 2-DG sensitivity, we next examined the responses of GFP-control and MAGEA6-expressing cells under glutamine-free conditions, thereby eliminating extracellular glutamine as a metabolic substrate (Fig. 4i-k). We generated 2-DG dose–response curves and evaluated cell viability using two complementary approaches. When viability was normalized to the corresponding untreated controls, MAGEA6-expressing cells exhibited a higher 2-DG IC50 than GFP-control cells, indicating greater relative resistance to glycolytic inhibition under glutamine deprivation (Fig. 4j). However, examination of absolute Alamar Blue fluorescence revealed a more complex, concentration-dependent response. At lower 2-DG concentrations, MAGEA6-expressing cells exhibited lower absolute viability than GFP controls, suggesting greater vulnerability to glutamine deprivation under conditions of limited glycolytic inhibition. Interestingly, this relationship reversed at higher 2-DG concentrations, where MAGEA6-expressing cells maintained greater viability than GFP controls (Fig. 4k). Thus, although MAGEA6-expressing cells exhibited greater vulnerability under glutamine-deprived conditions at lower levels of glycolytic stress, they gained a relative survival advantage as glycolytic inhibition intensified. These findings reveal a context-dependent metabolic response associated with MAGEA6 expression. Together, our data suggest that glutamine metabolism supports MIA PaCa-2 cell survival under 2-DG stress, as GLS1 inhibition increased sensitivity to it in both cell populations without abolishing the MAGEA6-associated resistance advantage. Complete glutamine deprivation further revealed that this advantage depends on the severity of glycolytic inhibition. Thus, MAGEA6-mediated resistance likely involves additional metabolic adaptations beyond increased reliance on glutamine metabolism.

Analysis of genes involved in glutamine metabolism also revealed distinct, condition-dependent transcriptional changes in MAGEA6-expressing cells (Fig. 4l). Under basal conditions, several genes involved in glutamine and amino acid metabolism, including GOT1, SLC38A4, and GLUD1, exhibited reduced expression relative to GFP controls. Interestingly, GOT1, encoding cytosolic aspartate aminotransferase, was downregulated in MAGEA6-expressing cells, whereas the transcript levels of its mitochondrial counterpart, GOT2, remained unchanged, suggesting differential regulation of these two enzymes involved in glutamine–aspartate metabolism [74]. Following 2-DG treatment, SLC1A3, encoding a glutamate/aspartate transporter, exhibited markedly increased expression in MAGEA6-expressing cells, particularly after 24 and 48 h. This finding raises the possibility that altered aspartate and glutamate transport may contribute to metabolic adaptation during glycolytic inhibition. In contrast, under glutamine-deprived conditions, GLS, ASNS, and PYCR2 exhibited increased expression in MAGEA6-expressing cells, suggesting additional adaptations involving glutamine metabolism, asparagine biosynthesis, and proline metabolism. Together, these findings indicate that MAGEA6 expression is associated with selective remodeling of amino acid transport and metabolism, with distinct transcriptional responses to glycolytic inhibition and glutamine deprivation.

### MAGEA6 discordantly regulates glycolytic enzyme levels and enhances cellular energy metabolism under 2-DG-induced metabolic stress

We next investigated whether MAGEA6 expression alters the abundance of key glycolytic enzymes, focusing initially on hexokinases (HK1 and HK2), which catalyze the first step of glycolysis and phosphorylate 2-DG (Fig. 5a). Western blot analysis following treatment with 3 mM 2-DG for 1, 6, or 24 h revealed reduced HK1 abundance in MAGEA6-expressing MIA PaCa-2 cells compared with GFP controls, particularly at later time points, while HK2 levels were lower after 24 h (Fig. 5b and S5a). Consistently, siRNA-mediated depletion of overexpressed or endogenous MAGEA in HeLa and DAOY cells increased HK1 protein abundance by approximately 1.6-fold and 2.6-fold, respectively (Fig. 5c and S5b–c). Reduced HK1 and HK2 abundance in MAGEA6-expressing cells was also observed following prolonged treatment with 2 mM 2-DG for 24 or 48 h, particularly at 24 h, whereas glutamine deprivation increased both HK isoforms in both cell populations (Fig. 5d–f). Interestingly, analysis of additional enzymes involved in glycolysis and pyruvate metabolism, including phosphofructokinase-1 (PFK1), pyruvate kinase M1/2 (PKM1/2), lactate dehydrogenase A (LDHA), and pyruvate dehydrogenase (PD), revealed increased abundance of several downstream enzymes in MAGEA6-expressing cells despite reduced HK1 and HK2 levels. Together, these findings reveal discordant regulation of glycolytic enzymes associated with MAGEA6 expression, characterized by reduced hexokinase abundance and increased levels of several downstream glycolytic enzymes under metabolic stress.

**Figure 5.**
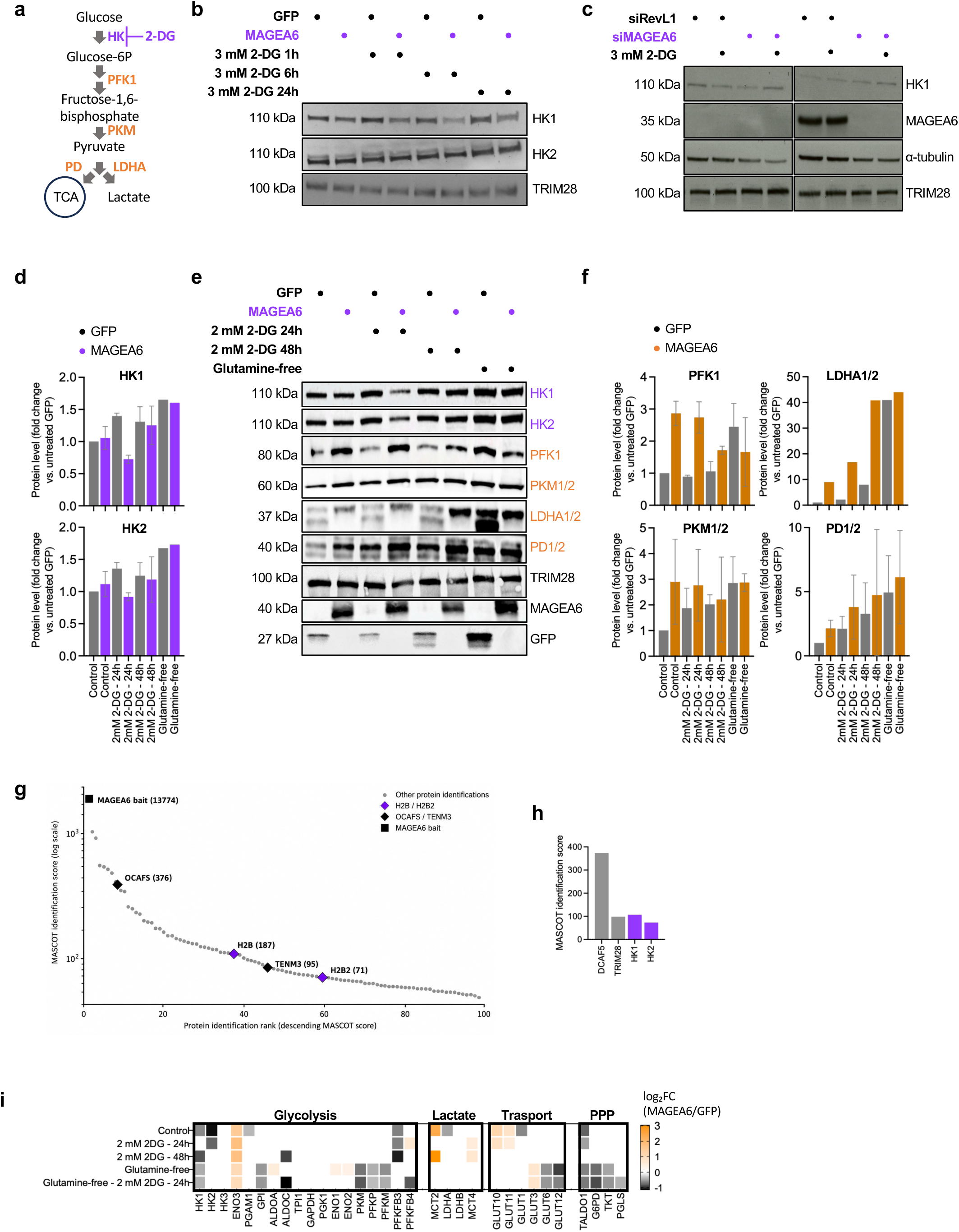
MAGEA6 differentially regulates glycolytic enzyme abundance under metabolic stress and is associated with hexokinases. **a.** Schematic representation of glycolysis and its major enzymes, including hexokinase (HK), phosphofructokinase-1 (PFK1), pyruvate kinase M (PKM), pyruvate dehydrogenase (PDH), and lactate dehydrogenase A (LDHA), highlighting key steps in glucose metabolism, pyruvate conversion to lactate, and pyruvate entry into the tricarboxylic acid (TCA) cycle. **b.** Representative Western blots showing HK1 and HK2 protein abundance in GFP-control and MAGEA6-expressing MIA PaCa-2 cells following treatment with 3 mM 2-deoxy-D-glucose (2-DG) for 1, 6, or 24 h. TRIM28 was used as a loading control. **c.** Representative Western blots showing HK1 and MAGEA6 protein abundance in MIA PaCa-2 cells transfected with MAGEA6-targeting or control siRNA and treated with 3 mM 2-DG. α-Tubulin and TRIM28 were used as loading controls. **d.** Densitometric quantification of HK1 and HK2 protein abundance in GFP-control and MAGEA6-expressing cells following treatment with 2 mM 2-DG for 24 or 48 h or culture in glutamine-free medium, based on the Western blots shown in panel e. **e.** Representative Western blots showing the abundance of glycolytic and associated metabolic enzymes, including HK1, HK2, PFK1, PKM1/2, LDHA1/2, and PD1/2, in GFP-control and MAGEA6-expressing cells under the indicated metabolic conditions. TRIM28 was used as a loading control, and GFP and MAGEA6 expression were confirmed by Western blotting. **f.** Densitometric quantification of PFK1, PKM1/2, LDHA1/2, and PD1/2 protein abundance, expressed as fold change relative to untreated GFP-control cells, based on the Western blots shown in panel e. **g–h.** Identification of candidate MAGEA6-associated proteins by tandem affinity purification coupled with mass spectrometry (TAP-MS) in HEK293 cells. **g.** Ranked dot plot showing 96 protein identification entries arranged in descending order of MASCOT identification score on a logarithmic scale. HK1 and HK2 are highlighted in purple; MAGEA6 (bait), TRIM28, and DCAF5 in black; and all remaining proteins in gray [22]. **h.** Bar graph comparing MASCOT identification scores of selected proteins, including MAGEA6 (bait), HK1, HK2, TRIM28, and DCAF5. HK1 and HK2 were identified with MASCOT scores of 107 and 73, respectively, compared with 98 for TRIM28 and 374 for DCAF5. Corresponding peptide counts and sequence coverage are provided in the accompanying table. MASCOT scores reflect protein identification evidence rather than relative protein abundance or interaction affinity. **i.** RNA-seq heatmap showing differential expression of genes involved in glucose metabolism and glycolytic regulation under the same experimental conditions. Genes are organized into four functional groups. Heatmap displays log₂ fold changes (MAGEA6/GFP) for genes meeting the differential-expression criteria (adjusted *P* ≤ 0.05 and |log₂FC| ≥ 1) in at least one experimental condition. Within each functional group, genes are ordered by descending log₂ fold change under untreated control conditions, with genes lacking reported control values positioned afterward. Data are presented as mean ± SD, where applicable.

To see whether MAGEA6 associates with glycolytic enzymes, we reanalyzed previously published tandem affinity purification–mass spectrometry (TAP-MS) data from HEK293 cells [75]. Among 96 protein identification entries, both HK1 and HK2 were identified as MAGEA6 co-purifying proteins, with MASCOT scores of 107 and 73, supported by 10 and 7 peptides and sequence coverage of 10.4% and 7.8%, respectively (Fig. 5g–h). HK1 and HK2 ranked 39th and 61st, respectively, with HK1 exhibiting a MASCOT score comparable to that of TRIM28, a known MAGEA-associated protein (score 98), whereas DCAF5 showed a substantially higher score (374). These findings suggest a previously uncharacterized potential association between MAGEA6 and hexokinases. In addition, transcriptomic analysis revealed a similar pattern of discordant glycolytic gene regulation, with reduced hexokinase expression alongside increased expression of several downstream glycolytic genes in MAGEA6-expressing cells (Fig. 5i). Together, these findings suggest that MAGEA6 modulates glycolytic metabolism at both transcriptional and protein levels, potentially through selective regulation of hexokinases rather than uniform suppression of glycolysis. We therefore investigated whether these changes were accompanied by alterations in cellular energy metabolism and mitochondrial function under metabolic stress.

### MAGEA6 alters cellular energy metabolism and lipid droplet accumulation under metabolic stress

To investigate whether MAGEA6-associated changes in glycolytic enzyme abundance affect cellular energy metabolism, we measured extracellular acidification rate (ECAR) and oxygen consumption rate (OCR) using the Seahorse XF Real-Time ATP Rate Assay (Fig. 6a). MAGEA6-expressing MIA PaCa-2 cells generally exhibited higher ECAR and OCR than GFP controls, particularly under 2-DG-induced metabolic stress, suggesting increased extracellular acidification and mitochondrial respiration. Oligomycin treatment increased ECAR while reducing OCR, consistent with compensatory glycolytic activity, whereas subsequent rotenone/antimycin A treatment further suppressed mitochondrial respiration. Together, these findings suggest that MAGEA6-mediated resistance to glycolytic stress is associated with altered energy metabolism despite reduced hexokinase abundance.

**Figure 6.**
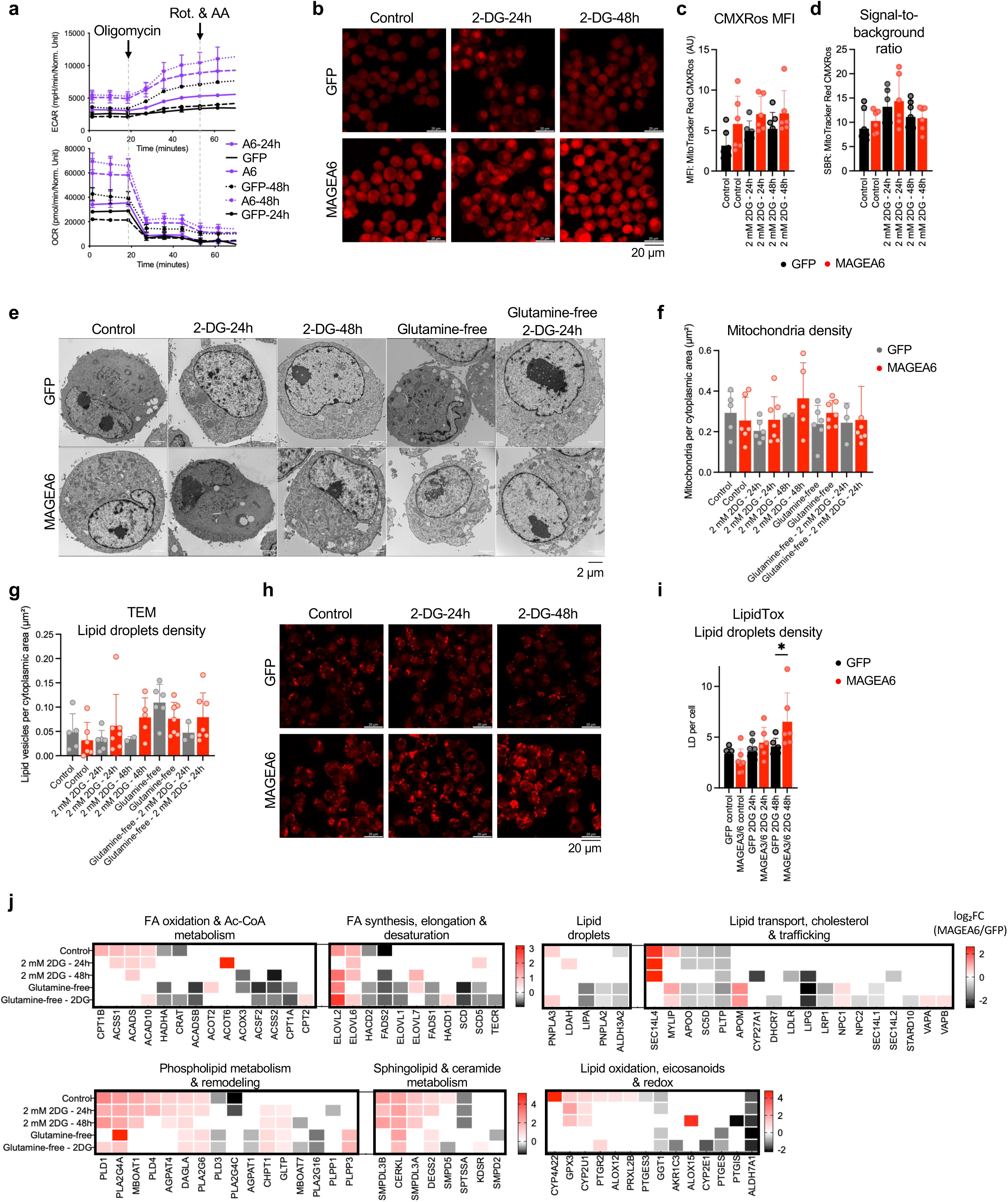
MAGEA6 expression alters cellular energy metabolism, mitochondrial membrane potential-associated fluorescence, lipid droplet accumulation, and lipid metabolic gene expression in pancreatic cancer cells. **a.** Seahorse real-time ATP rate assay measuring extracellular acidification rate (ECAR, upper panel) and oxygen consumption rate (OCR, lower panel) in GFP-control and MAGEA6-expressing MIA PaCa-2 cells under the indicated metabolic conditions. Oligomycin and rotenone/antimycin A were sequentially injected to assess glycolytic and mitochondrial contributions to cellular ATP production. **b.** Representative confocal fluorescence microscopy images of GFP-control and MAGEA6-expressing MIA PaCa-2 cells stained with MitoTracker Red CMXRos under untreated conditions and following 24 or 48 h of treatment with 2 mM 2DG. CMXRos fluorescence was used to assess mitochondrial membrane potential-dependent dye accumulation. Scale bars are indicated. **c.** Quantification of MitoTracker Red CMXRos mean fluorescence intensity (MFI), expressed in arbitrary units (AU), in GFP-control and MAGEA6-expressing cells under the indicated treatment conditions. **d.** Quantification of MitoTracker Red CMXRos signal-to-background ratio (SBR), calculated as the ratio of cellular fluorescence intensity to background fluorescence intensity under the corresponding experimental conditions. **e.** Representative transmission electron microscopy (TEM) images showing cellular ultrastructure, mitochondrial morphology, and lipid droplets in GFP-control and MAGEA6-expressing MIA PaCa-2 cells under the indicated metabolic conditions. Scale bars are indicated. **f.** Quantification of mitochondrial area relative to the analyzed cellular area in TEM images of GFP-control and MAGEA6-expressing cells under the indicated metabolic conditions. **g.** Quantification of lipid droplet density, expressed as the number of lipid droplets per unit cytoplasmic area, in TEM images of GFP-control and MAGEA6-expressing cells under the indicated metabolic conditions. **h.** Representative confocal fluorescence microscopy images of GFP-control and MAGEA6-expressing MIA PaCa-2 cells stained with HCS LipidTOX Deep Red Neutral Lipid Stain under untreated conditions and following 24 or 48 h of treatment with 2 mM 2DG. Scale bars are indicated. **i.** Quantification of lipid droplet number per cell in GFP-control and MAGEA6-expressing cells under the indicated treatment conditions. Lipid droplets were identified by fluorescence thresholding and particle analysis using Fiji/ImageJ, and the total number of lipid droplets was normalized to the number of DAPI-positive nuclei in each image. **j.** Heatmaps showing log2 fold changes (MAGEA6/GFP) in the expression of genes involved in lipid metabolism, grouped by their primary biological functions. Rows represent five metabolic conditions. Genes were selected based on significant differential expression in at least one experimental condition (adjusted P < 0.05) and organized according to their primary metabolic functions. Positive and negative values indicate increased and decreased expression, respectively, in MAGEA6-expressing cells relative to GFP controls. Gray indicates values not displayed for the corresponding comparison. Data are presented as mean ± SD, where applicable. Statistical comparisons in panels c, d, f, g, and i were performed using one-way ANOVA followed by Šidák’s multiple-comparisons test. *P* < 0.05 was considered statistically significant.

To further assess mitochondrial responses, we stained GFP-control and MAGEA6-expressing cells with MitoTracker Red CMXRos, a membrane potential-dependent fluorescent dye (Fig. 6b–d). Quantification of mean fluorescence intensity (MFI) and signal-to-background ratio (SBR) revealed variability across experimental conditions (Fig. 6c–d). Although one-way ANOVA identified a significant overall effect on MFI (*P* = 0.039), individual pairwise comparisons were not significant after correction for multiple testing. Nevertheless, MAGEA6-expressing cells generally showed a trend toward higher CMXRos fluorescence, suggesting potential differences in mitochondrial membrane potential and/or abundance. We next examined mitochondrial content and lipid storage using transmission electron microscopy (TEM; Fig. 6e-g). Quantification revealed nonsignificant trends toward greater mitochondrial area and lipid droplet density in MAGEA6-expressing cells following 2-DG treatment (Fig. 6f–g). Complementary LipidTOX Deep Red staining further supported increased neutral lipid droplet accumulation in MAGEA6-expressing cells under prolonged glycolytic stress (Fig. 6h–i).

To investigate the effects of MAGEA6 on lipid metabolism, we compared lipid metabolism-related gene expression between MAGEA6-expressing and GFP-control cells across five metabolic conditions (Fig. 6j). MAGEA6 expression was associated with higher levels of genes involved in phospholipid remodeling (*PLA2G4A*, *PLD1*, *DAGLA*), fatty acid elongation (*ELOVL2*, *ELOVL6*), and sphingolipid metabolism (*CERKL*, *DEGS2*) across multiple conditions, whereas *ALDH7A1*, *PLTP*, and *FADS2* were consistently expressed at lower levels. MAGEA6 differentially affected fatty acid oxidation-associated genes, with higher *CPT1B*, *ACADS*, and *ACSS1* expression under basal conditions but lower *CPT1A*, *HADHA*, and *ACADSB* expression during glutamine deprivation. MAGEA6 also influenced lipid droplet-associated genes, including higher *PNPLA3* and *LDAH* expression under selected conditions, whereas *PNPLA2* and *LIPA*, involved in triglyceride and lysosomal lipid hydrolysis, respectively, showed lower expression during glutamine deprivation. These MAGEA6-associated transcriptional differences, together with increased lipid droplet accumulation following 2-DG treatment, suggest that MAGEA6 influences lipid storage, remodeling, and metabolic adaptation under glycolytic stress.

### Integrated multi-omics analysis reveals MAGEA3/6-associated metabolic adaptations to glycolytic inhibition and glutamine deprivation and identifies a potential therapeutic vulnerability

To determine whether these transcriptional differences were accompanied by alterations in protein abundance, we performed comparative proteomic analyses across the same five experimental conditions (Fig. 7a-e). Under basal conditions (Fig. 7a), MAGEA6-expressing cells exhibited increased abundance of several proteins involved in mitochondrial function and cellular stress responses, including GRPE1, NDUFA2, HIGD1A, and COQ9 [76–79]. Transglutaminase 2 (TGM2), a multifunctional protein implicated in cellular stress responses and survival signaling, was approximately 10-fold more abundant in MAGEA6-expressing cells than in GFP controls (log2 fold change = 3.30). In contrast, PCYT1B, an enzyme involved in phosphatidylcholine biosynthesis through the Kennedy pathway, was substantially reduced (log2 fold change = −1.67), suggesting differences in membrane phospholipid metabolism even under basal conditions [80, 81]. Following 24 h of 2-DG, proteomic analysis revealed increased abundance of several mitochondrial metabolic enzymes in MAGEA6-expressing cells, including dihydrolipoamide S-succinyltransferase (DLST; log2 fold change = 2.86), serine hydroxymethyltransferase 2 (SHMT2; 1.65), succinate dehydrogenase subunit B (SDHB; 1.57), mitochondrial aspartate aminotransferase (GOT2; 1.41), and mitochondrial malate dehydrogenase (MDH2; 1.30) (Fig. 7b). These proteins participate in the tricarboxylic acid (TCA) cycle, mitochondrial respiration, glutamine–aspartate metabolism, and one-carbon metabolism, suggesting that MAGEA6 expression is associated with coordinated alterations in mitochondrial metabolic pathways during glycolytic inhibition. Particularly interesting was the increased abundance of mitochondrial GOT2 protein, contrasting with the downregulation of cytosolic *GOT1* at the transcriptional level observed in our RNA-seq analysis. Together, these findings suggest differential regulation of mitochondrial and cytosolic aspartate aminotransferases in MAGEA6-expressing cells, potentially reflecting altered glutamine–aspartate metabolism and cellular redox homeostasis.

**Figure 7.**
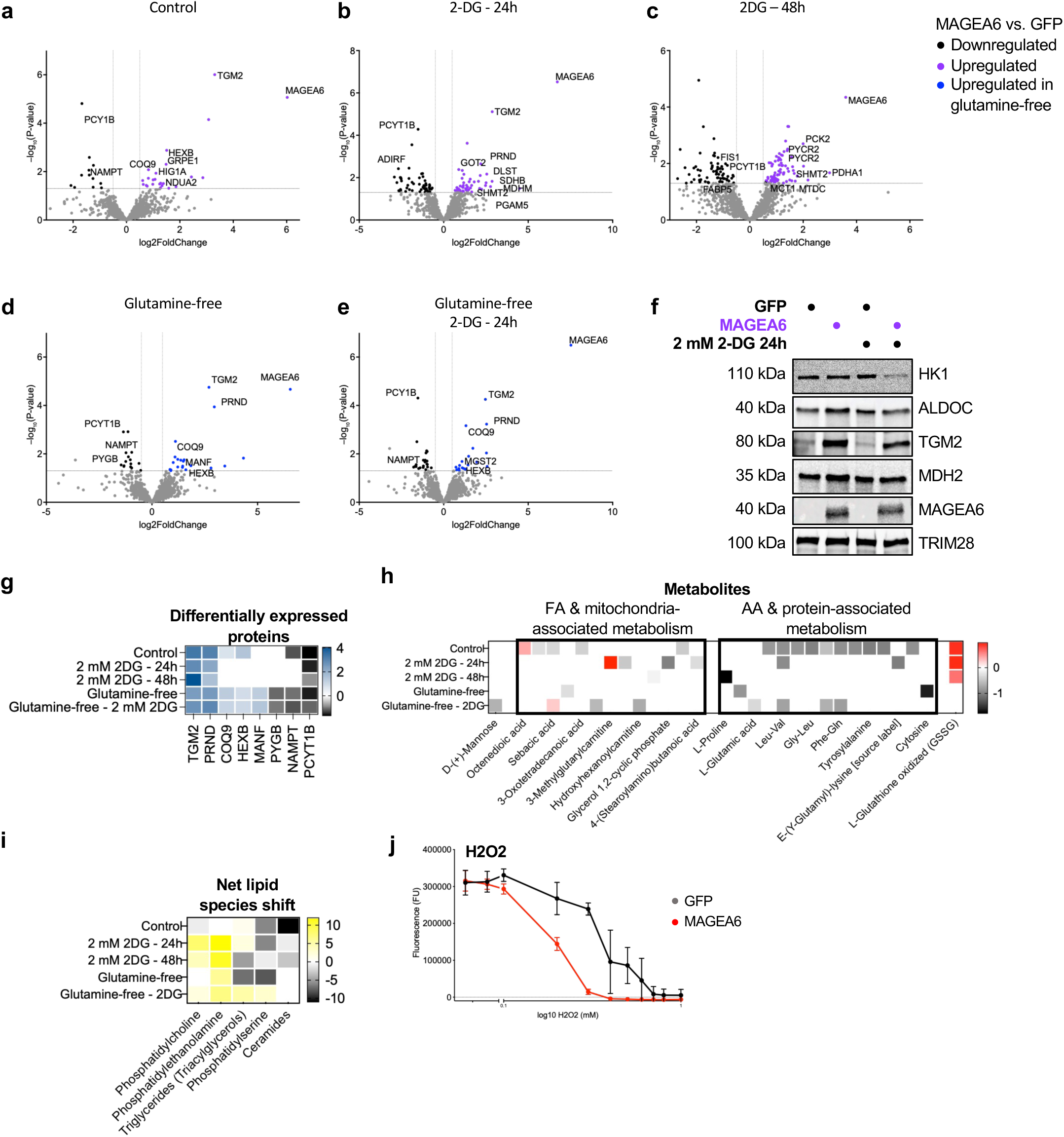
Integrated proteomic, metabolomic, and lipidomic profiling reveals MAGEA6-associated metabolic adaptations under 2-DG-induced glycolytic inhibition and glutamine deprivation and identifies a potential vulnerability to oxidative stress. a–e. Volcano plots showing differential protein abundance between MAGEA6-expressing and GFP-control MIA PaCa-2 cells under **a.** untreated control conditions, **b.** treatment with 2 mM 2-DG for 24 h, **c.** treatment with 2 mM 2-DG for 48 h, **d.** glutamine-free conditions, and **e.** glutamine-free conditions combined with 2 mM 2-DG treatment for 24 h. Each point represents an identified protein, plotted according to its log₂ fold change (MAGEA6/GFP) and −log₁₀(*P*-value). Differentially abundant proteins are highlighted in purple or blue (increased abundance) and black (decreased abundance), whereas gray points represent proteins that do not meet the indicated significance threshold. Selected proteins involved in mitochondrial metabolism, cellular stress responses, and lipid homeostasis are labeled.**f.** Representative Western blots showing hexokinase 1 (HK1), aldolase C (ALDOC), transglutaminase 2 (TGM2), and mitochondrial malate dehydrogenase 2 (MDH2) protein abundance in GFP-control and MAGEA6-expressing MIA PaCa-2 cells under untreated conditions and following treatment with 2 mM 2-DG for 24 h. MAGEA6 expression was confirmed by Western blotting, and TRIM28 was used as a loading reference. **g.** Heatmap showing shared and condition-specific differences in protein abundance between MAGEA6-expressing and GFP-control cells across the five experimental conditions. Values represent log₂ fold changes (MAGEA6/GFP) for selected proteins involved in cellular stress responses and metabolic regulation. Purple indicates increased protein abundance, whereas dark gray to black indicates decreased abundance in MAGEA6-expressing cells. White indicates proteins not identified among the differentially abundant proteins in the corresponding comparison. **h.** Heatmap showing differential metabolite abundance between MAGEA6-expressing and GFP-control MIA PaCa-2 cells across the five experimental conditions. Selected metabolites involved in carbohydrate, amino acid, lipid, and redox metabolism are shown. Values represent log₂ fold changes (MAGEA6/GFP), with red indicating increased abundance and dark gray to black indicating decreased abundance. **i.** Heatmap summarizing MAGEA6-associated changes in major lipid classes across the five experimental conditions. Values represent the net number of differential lipid features, calculated as the number of features increased minus the number decreased in MAGEA6-expressing cells relative to GFP controls within each condition. Counts represent lipid features reported in the source comparisons rather than total lipid class abundance or FDR-adjusted significance. **j.** Dose–response analysis of GFP-control and MAGEA6-expressing MIA PaCa-2 cells following treatment with increasing concentrations of hydrogen peroxide (H₂O₂). Relative cell viability was assessed using the AlamarBlue assay.

Following 48 h of 2-DG treatment, several additional mitochondrial metabolic proteins exhibited higher abundance in MAGEA6-expressing cells, including pyruvate dehydrogenase E1 subunit alpha 1 (PDHA1; log2 fold change = 2.99), mitochondrial phosphoenolpyruvate carboxykinase (PCK2; 2.00), mitochondrial methylenetetrahydrofolate dehydrogenase (MTHFD2; 1.78), and serine hydroxymethyltransferase 2 (SHMT2; 1.66) (Fig. 7c). These enzymes participate in mitochondrial carbon metabolism, pyruvate utilization, and one-carbon metabolism, suggesting that MAGEA6 expression is associated with broader metabolic remodeling during prolonged glycolytic inhibition. These proteomic findings complement our Seahorse analyses showing altered mitochondrial respiratory activity in MAGEA6-expressing cells, supporting a potential role for mitochondrial metabolic adaptation in resistance to glycolytic stress, although changes in metabolic flux remain to be established [82]. Under glutamine-deprived conditions, MAGEA6-expressing cells had increased levels of proteins associated with mitochondrial electron transport and homeostasis, including NDUFA3 (2.53), COQ9 (1.32), and MITOK (1.44), as well as proteins associated with cellular stress responses and lipid metabolism, including MANF, MGST2, and HEXB (Fig. 7d). In contrast, reduced level was observed under combined glutamine deprivation and 2-DG treatment, with increased abundance of NDUFC2, MANF, COQ9, and HEXB and reduced abundance of NAMPT, PYGB, and ALDH3A2 (Fig. 7e). These findings suggest that MAGEA6 expression is associated with alterations in mitochondrial homeostasis, cellular stress responses, and lipid and carbohydrate metabolism under nutrient-limited conditions.

Across all five experimental conditions, two proteins exhibited particularly consistent differences between MAGEA6-expressing and GFP-control cells. TGM2 remained strongly elevated, with log2 fold changes ranging from 2.47 to 4.05, whereas PCYT1B remained reduced, with log2 fold changes ranging from −0.84 to −1.67 (Fig.7g). Additional proteins, including COQ9, HEXB, NAMPT, and PYGB, exhibited recurring differences in abundance across several metabolic conditions, further supporting alterations in mitochondrial function, phospholipid metabolism, and cellular metabolic homeostasis. Representative Western blot analysis of selected metabolic and stress-associated proteins, including HK1, ALDOC, TGM2, and MDH2, was performed to further examine protein abundance in GFP-control and MAGEA6-expressing cells under basal conditions and following 2-DG treatment (Fig. 7f). Collectively, integrated transcriptomic and proteomic analyses reveal that MAGEA6 expression is associated with broad but condition-dependent metabolic reprogramming involving mitochondrial carbon metabolism, glutamine–aspartate metabolism, cellular stress responses, and lipid homeostasis.

To further characterize MAGEA6-associated metabolic adaptations, we analyzed metabolite abundance across five experimental conditions (Fig. 8h). Metabolomic profiling revealed distinct differences in carbohydrate, amino acid, lipid, and redox metabolism between MAGEA6-expressing and GFP-control cells. In carbohydrate and nucleotide metabolism, MAGEA6-expressing cells exhibited reduced D-mannose abundance under combined glutamine deprivation and 2-DG treatment and lower cytosine levels during glutamine deprivation, complementing our observations of discordant glycolytic enzyme regulation, particularly reduced HK1 and HK2 protein abundance despite increased levels of several downstream enzymes. Amino acid and peptide metabolism was also altered, with reduced abundance of several dipeptides, including Leu-Val, Gly-Leu, and Phe-Gln. Interestingly, Leu-Val remained lower across multiple metabolic conditions, while proline abundance decreased following 48 h of 2-DG treatment despite increased abundance of the proline biosynthesis enzymes PYCR1 and PYCR2. Additional differences included reduced serine and glutamate under glutamine-deprived conditions. Proteomic analysis further revealed increased abundance of mitochondrial metabolic enzymes (Fig. 7a-g), including DLST, SDHB, and MDH2, following 24 h of 2-DG treatment, consistent with transcriptomic evidence of altered glutamine–aspartate metabolism and increased mitochondrial oxygen consumption.

Lipid metabolism also exhibited MAGEA6-associated changes, including differences in dicarboxylic fatty acids, acylcarnitines, and lipid-derived metabolites. Following 24h of 2-DG, 3-methylglutarylcarnitine was increased, whereas 2-hexenoylcarnitine was reduced; hydroxyhexanoylcarnitine was also reduced under combined glutamine deprivation and 2-DG treatment. These findings complement differential expression of fatty acid metabolism genes (*CPT1B*, *CPT1A*, *ACADS*, *HADHA*) (Fig. 6j) and lipid remodeling genes, together with reduced PCYT1B protein abundance. Interestingly, *ALDH3A2* showed concordant reductions at both transcript and protein levels under combined metabolic stress, further supporting altered lipid metabolic regulation. Finally, oxidized glutathione (GSSG) was consistently elevated in MAGEA6-expressing cells under basal and 2-DG-treated conditions, suggesting altered redox homeostasis, potentially associated with changes in mitochondrial metabolism and increased *GPX3* expression. Together, these findings indicate that MAGEA6-associated resistance to glycolytic stress coincides with broad metabolic adaptations involving mitochondrial carbon metabolism, amino acid homeostasis, lipid remodeling, and redox regulation, extending beyond the discordant regulation of glycolytic enzymes.

To further investigate MAGEA6-associated lipid metabolic adaptations, we performed lipidomic profiling of MAGEA6-expressing and GFP-control MIA PaCa-2 cells across five metabolic conditions (Fig. S3 and 8i). Lipidomic analysis revealed extensive remodeling of membrane phospholipids, sphingolipids, and triglycerides associated with MAGEA6 expression. Notably, phosphatidylethanolamine (PE) species were consistently more abundant in MAGEA6-expressing cells following 2-DG treatment, with all 12 identified differential PE features increased at 24 h and all 11 increased at 48 h. Several phosphatidylcholine (PC) species, particularly ether-linked PC species, were also elevated, suggesting altered membrane phospholipid composition during glycolytic stress. These findings complement transcriptomic changes in phospholipid remodeling genes, including *PLA2G4A*, *PLD1*, and *CHPT1*, together with reduced abundance of the phosphatidylcholine biosynthesis enzyme PCYT1B identified by proteomics.

Sphingolipid metabolism was also altered, with most differential ceramide species exhibiting lower abundance in MAGEA6-expressing cells under basal conditions and following prolonged 2-DG treatment. These observations are consistent with broader changes in sphingolipid-associated gene expression, including *CERKL* and *DEGS2*, and suggest altered sphingolipid homeostasis. Triglycerides exhibited more heterogeneous, condition-dependent differences, with predominantly lower levels under glutamine deprivation and at 48 h of 2-DG treatment, whereas most differential triglyceride species were elevated under combined glutamine deprivation and 2-DG exposure. These changes complement our observations of altered lipid droplet accumulation and differential expression of lipid storage and lipolysis-associated genes, including *PNPLA3*, *PNPLA2*, and *LIPA*.

Interestingly, lipidomic analysis revealed MAGEA6-associated remodeling of lipid species potentially involved in oxidative stress responses. Ether-linked phosphatidylethanolamine (PE) species, including plasmalogens, were enriched in MAGEA6-expressing cells following 2-DG treatment, whereas several ceramide species exhibited reduced abundance. These changes, together with elevated oxidized glutathione (GSSG), increased *GPX3* expression, and altered mitochondrial metabolism, suggest that MAGEA6 influences cellular redox homeostasis. Remarkably, despite their increased resistance to 2-DG-induced glycolytic stress, MAGEA6-expressing cells exhibited greater sensitivity to hydrogen peroxide (H₂O₂)-induced oxidative stress (Fig. 7j), revealing a potential metabolic vulnerability associated with MAGEA6 expression. Together, these data demonstrate that MAGEA6 promotes extensive remodeling of membrane phospholipids, sphingolipids, and neutral lipids during metabolic stress. The enrichment of PE species, altered mitochondrial metabolism, and increased lipid droplet accumulation suggest that lipid metabolic adaptations may contribute to MAGEA6-mediated resistance to glycolytic inhibition. Importantly, the increased susceptibility to oxidative stress indicates that this protective phenotype is stress-specific and may expose a potential therapeutic vulnerability in MAGEA6-expressing pancreatic cancer cells.

## DISCUSSION

In the present work, we investigated the role of MAGEA3/6 in metabolic plasticity and resistance to nutrient deprivation and metabolic stress in pancreatic cancer cells, building on our previous finding that MAGEA3/6 confer resistance to 2-deoxyglucose (2-DG)-mediated glycolytic inhibition [14, 83]. Our objective was to elucidate the molecular mechanisms underlying this resistance and determine whether MAGEA3/6 promote broader metabolic adaptations that support pancreatic cancer cell survival under nutrient-limited conditions. Metabolic plasticity is a hallmark of PDAC, enabling cancer cells to adapt to nutrient limitation through changes in glycolysis, mitochondrial metabolism, and utilization of alternative metabolic substrates [84]. Building on our previous findings [28], we showed that MAGEA3/6 expression enhances resistance to 2-DG in highly glycolytic MIA PaCa-2 cells. Although cell viability decreased with increasing 2-DG concentrations, MAGEA3/6-expressing cells consistently exhibited greater resistance than GFP controls. A similar phenotype was observed in PANC-1 cells, indicating that this protective effect is not restricted to a single PDAC model. These findings are consistent with the established dependence of pancreatic cancer cells on glycolytic metabolism and their vulnerability to its inhibition [85–88]. Interestingly, MAGEA3/6-mediated resistance was specific to certain metabolic challenges. Unlike 2-DG, treatment with 3-bromopyruvate (3-BP) did not reveal significant differences in sensitivity between MAGEA3/6-expressing cells and controls. Because 3-BP affects multiple metabolic processes, including glycolysis, mitochondrial function, and cellular redox homeostasis [89, 90], its broader metabolic effects may overcome the protective advantage associated with MAGEA3/6 expression. Similarly, inhibition of the pentose phosphate pathway (PPP) with 6-aminonicotinamide (6-AN) increased sensitivity to glycolytic stress but did not eliminate the relative resistance of MAGEA3/6-expressing cells. Although increased PPP activity has been implicated in pancreatic cancer resistance to oxidative stress and radiation [91], our findings suggest that MAGEA3/6-mediated resistance to 2-DG cannot be explained only by increased reliance on the PPP.

Previous studies identified MAGEA3/6 as regulators of AMPK signaling through their interaction with TRIM28, promoting AMPK ubiquitination and degradation, reducing autophagy, and enhancing mTOR signaling [33, 92]. Pharmacological activation of AMPK has also been shown to suppress MAGEA6-associated oncogenic phenotypes [16]. However, our results did not show the expected reduction in AMPKα1 protein abundance in MAGEA6-expressing cells. Instead, increased expression of AMPK-associated transcripts, particularly *PRKAA2* (AMPKα2), as well as increased AMPK protein abundance, was observed under selected experimental conditions, suggesting that MAGEA6-mediated regulation of AMPK may be context-dependent and occur at both transcriptional and protein levels. Moreover, differences in LC3 abundance and the response to mTORC1 inhibition with rapamycin did not support a simple model in which AMPK degradation and mTOR activation fully explain MAGEA6-mediated resistance to glycolytic inhibition. Although rapamycin enhanced sensitivity to 2-DG, MAGEA6-expressing cells maintained a relative resistance advantage, suggesting that additional metabolic pathways contribute to this protective phenotype.

One of the most striking findings was the discordant regulation of glycolytic enzymes in MAGEA6-expressing cells. HK1 and HK2 exhibited reduced abundance, whereas several downstream glycolytic enzymes were increased. Transcriptomic analysis further revealed selective regulation of glycolytic genes, suggesting that MAGEA6 influences glycolysis through both transcriptional and protein-level mechanisms.

Hexokinases catalyze the first step of glucose metabolism, and their activity is influenced not only by protein abundance but also by post-translational modifications and mitochondrial localization. Previous studies demonstrated that ubiquitination and SUMOylation can regulate HK2 mitochondrial association, glycolytic activity, and cancer cell survival [93]. Other studies have linked altered expression of hexokinase family members to tumor progression [94]. Furthermore, previous affinity purification–mass spectrometry analysis of MAGEA6 expressed in HEK293 cells identified both HK1 and HK2 among the co-purifying proteins, alongside the previously established MAGEA3/6-interacting partner TRIM28. These findings suggest a potential association between MAGEA6 and hexokinases, raising the possibility that MAGEA6 influences glycolytic metabolism through interactions with HK1 and HK2, in addition to transcriptional regulation [95]. However, whether these associations are direct and contribute to hexokinase stability, activity, or subcellular localization remains to be determined. Furthermore, our transcriptomic data suggest that reduced HK1 and HK2 abundance in MAGEA6-expressing cells may also involve transcriptional regulation. Together, these findings suggest that MAGEA6 influences hexokinase expression through multiple regulatory mechanisms, potentially contributing to selective remodeling of glycolytic metabolism rather than a uniform reduction in glycolytic capacity.

Interestingly, reduced HK1 and HK2 abundance in MAGEA6-expressing cells was accompanied by increased expression of several downstream glycolytic enzymes, revealing discordant regulation of glycolysis rather than uniform suppression of the pathway and suggesting selective metabolic reprogramming despite reduced hexokinase abundance. Consistent with this observation, Seahorse analyses demonstrated higher oxygen consumption rates (OCR) and extracellular acidification rates (ECAR) in MAGEA6-expressing cells, particularly following 2-DG treatment, indicating increased mitochondrial respiratory activity and extracellular acidification. MitoTracker analysis also revealed a trend toward increased mitochondrial staining, although individual group comparisons were not statistically significant. Furthermore, integrated transcriptomic and proteomic analyses identified increased abundance of several mitochondrial enzymes involved in the TCA cycle, pyruvate metabolism, and amino acid utilization, including DLST, SDHB, MDH2, PDHA1, PCK2, SHMT2, and MTHFD2. Together, these findings suggest that MAGEA3/6-mediated resistance to glycolytic inhibition is associated with coordinated remodeling of glycolytic and mitochondrial metabolic pathways, potentially enhancing metabolic flexibility and supporting pancreatic cancer cell survival under metabolic stress despite reduced hexokinase abundance [96].

Glutamine metabolism may provide an additional mechanism supporting metabolic adaptation. Glutamine supplies carbon and nitrogen for mitochondrial metabolism, biosynthetic processes, and redox homeostasis, particularly when glucose utilization is limited [97, 98]. Interestingly, *GOT1*, encoding cytosolic aspartate aminotransferase, was transcriptionally downregulated in MAGEA6-expressing cells, whereas *GOT2* transcript abundance remained largely unchanged. In contrast, mitochondrial GOT2 protein abundance was increased following 24 h of 2-DG treatment. GOT1 and GOT2 participate in the noncanonical glutamine metabolic pathway characteristic of KRAS-driven PDAC, which supports NADPH production and cellular redox balance [74]. These findings suggest differential regulation of mitochondrial and cytosolic aspartate metabolism, potentially influencing glutamine utilization during glycolytic stress. Furthermore, although glutaminase inhibition increased sensitivity to 2-DG, MAGEA3/6-expressing cells retained a relative resistance advantage, suggesting that their metabolic adaptation is not entirely dependent on glutaminase activity. Complete glutamine deprivation further revealed that this advantage depends on the severity of glycolytic stress, highlighting the context-dependent nature of MAGEA6-mediated metabolic protection.

Alterations in lipid metabolism may also contribute to this phenotype. Our earlier observations of increased fatty acid oxidation associated with MAGEA6 expression [14, 83], together with the present findings of altered lipid droplet accumulation under metabolic stress, suggest changes in lipid metabolic regulation. Transcriptomic analysis revealed heterogeneous regulation of lipid metabolism-related genes, while proteomic analysis consistently identified reduced abundance of PCYT1B, an enzyme involved in phosphatidylcholine biosynthesis. Because phosphatidylcholine is an essential component of cellular membranes and lipid droplet surfaces, altered PCYT1B abundance may reflect changes in phospholipid homeostasis [99]. Additionally, transglutaminase 2 (TGM2), a multifunctional protein implicated in cellular stress responses and survival signaling, was consistently elevated in MAGEA6-expressing cells across all metabolic conditions. These findings suggest that lipid remodeling and stress-response pathways may contribute to MAGEA6-associated metabolic plasticity, although their direct involvement in resistance to glycolytic inhibition remains to be established [80].

Our integrated multi-omics analyses further support a broader role for MAGEA6 in metabolic reprogramming. Proteomic profiling revealed increased abundance of mitochondrial enzymes involved in carbon metabolism, amino acid utilization, and cellular stress responses, whereas metabolomic analysis identified alterations in amino acid and peptide metabolism, fatty acid derivatives, and redox homeostasis. Interestingly, reduced proline abundance following prolonged 2-DG treatment was accompanied by increased abundance of the proline biosynthesis enzymes PYCR1 and PYCR2, suggesting altered proline homeostasis. Lipidomic profiling revealed extensive remodeling of membrane phospholipids, sphingolipids, and triglycerides, with particularly pronounced enrichment of phosphatidylethanolamine (PE) species following 2-DG treatment. Increased PE abundance may reflect adaptations in membrane lipid composition relevant to mitochondrial function, whereas reduced ceramide abundance may be associated with altered cellular stress signaling. In contrast, heterogeneous changes in triglyceride species suggest selective lipid remodeling rather than uniform lipid accumulation. These findings complement transcriptomic changes in lipid remodeling and metabolism-related genes, including *PLA2G4A*, *PLD1*, *ELOVL2*, *PNPLA3*, and *LIPA*, together with reduced PCYT1B protein abundance and altered lipid droplet accumulation. Although these observations do not directly establish changes in mitochondrial membrane function, ceramide-mediated stress signaling, or fatty acid oxidation, they suggest that MAGEA6 promotes context-dependent lipid metabolic remodeling that may contribute to cellular adaptation during glycolytic stress.

Previous studies demonstrated that 2-DG and glucose deprivation can induce oxidative stress in pancreatic cancer cells, accompanied by increased oxidized glutathione (GSSG) levels and altered NADP⁺/NADPH ratios [47]. Consistent with these observations, our metabolomic analysis revealed elevated GSSG in MAGEA6-expressing cells under basal conditions and following 2-DG treatment, suggesting altered glutathione redox homeostasis. Furthermore, lipidomic analysis identified increased abundance of ether-linked phospholipids, including plasmalogens, and reduced abundance of several ceramide species, potentially reflecting changes in membrane lipid composition associated with oxidative stress responses. Together with increased *GPX3* expression and altered mitochondrial metabolism, these observations suggest that MAGEA6-associated metabolic adaptation may involve changes in cellular redox regulation. However, increased GSSG and altered lipid composition alone do not establish enhanced antioxidant protection or increased lipid peroxidation.

In all, our study identifies MAGEA3/6 as potential regulators of metabolic plasticity in PDAC. Rather than acting exclusively through the previously described AMPK–mTOR pathway, MAGEA3/6-associated resistance to glycolytic inhibition involves selective regulation of glycolytic enzymes and broader adaptations in mitochondrial carbon metabolism, glutamine–aspartate metabolism, lipid homeostasis, and cellular stress responses. These adaptations may enhance the ability of pancreatic cancer cells to tolerate metabolic challenges, highlighting MAGEA3/6-associated pathways as potential targets for further mechanistic investigation. Importantly, understanding the molecular functions of MAGEA3/6 and their contributions to metabolic reprogramming may also reveal previously unrecognized therapeutic vulnerabilities. Indeed, in striking contrast to their enhanced resistance to 2-DG-induced glycolytic stress, MAGEA6-expressing cells exhibited greater sensitivity to hydrogen peroxide (H₂O₂)-induced oxidative stress. This finding suggests that MAGEA6-mediated metabolic adaptations confer selective protection rather than broad resistance to cellular stress and may create distinct metabolic liabilities. Thus, elucidating the mechanisms by which MAGEA3/6 regulate cancer metabolism may not only improve our understanding of metabolic plasticity and therapy resistance but also uncover new opportunities to selectively target MAGEA3/6-expressing pancreatic cancer cells.

## LIMITATIONS OF THE STUDY

One limitation of this study is the use of the AlamarBlue assay to assess cell viability, as it measures cellular metabolic reducing activity rather than directly quantifying viable cell numbers. This distinction is particularly relevant given the metabolic reprogramming associated with MAGEA3/6 expression. Therefore, changes in AlamarBlue fluorescence should be interpreted as differences in relative cell viability and metabolic activity rather than exclusively reflecting cell survival. However, our findings were further supported by complementary assays assessing cell growth and clonogenic capacity.

Another limitation is that our findings were primarily obtained using *in vitro* cancer cell models, and the contribution of MAGEA3/6 to metabolic adaptation and resistance to glycolytic inhibition remains to be validated *in vivo*. Furthermore, although our integrated transcriptomic, proteomic, metabolomic, and lipidomic analyses identified extensive MAGEA6-associated metabolic changes, these approaches primarily measure molecular abundance rather than metabolic pathway activity. Direct metabolic flux analyses, including isotope tracing, would be needed to establish whether the observed changes reflect altered glycolytic, glutamine, mitochondrial, or lipid metabolic fluxes.

Although previous studies established interactions between MAGEA3/6 and TRIM28, and our reanalysis of affinity purification–mass spectrometry data identified HK1 and HK2 as potential MAGEA6-associated proteins, further validation by co-immunoprecipitation and complementary protein interaction assays is needed to confirm these associations in pancreatic cancer cells. Additional studies are also required to determine whether MAGEA6 regulates hexokinase abundance through transcriptional mechanisms, post-transcriptional regulation, or changes in protein stability and degradation, and whether these mechanisms influence hexokinase enzymatic activity, post-translational modifications, or subcellular localization.

Finally, although altered glutathione and lipid metabolism, together with increased sensitivity to oxidative stress, suggest potential vulnerabilities associated with MAGEA6 expression, the underlying mechanisms and their therapeutic relevance require further investigation. Despite these limitations, our findings provide new insights into the role of MAGEA3/6 in metabolic plasticity and identify promising directions for future mechanistic and therapeutic studies.

## CONCLUSIONS

Our findings identify MAGEA3/6 as regulators of metabolic plasticity, extending their previously established protective role in spermatogenesis during nutritional stress to pancreatic cancer cells. MAGEA3/6 promote resistance to 2-DG-induced glycolytic stress through metabolic adaptations extending beyond the previously described AMPK–mTOR signaling pathway. Integrated multi-omics analyses revealed selective regulation of glycolytic enzymes and extensive remodeling of mitochondrial metabolism, glutamine utilization, lipid homeostasis, and cellular stress responses. Importantly, despite enhanced resistance to glycolytic inhibition, MAGEA6-expressing cells exhibited increased sensitivity to oxidative stress. Given the particular vulnerability of spermatogenesis to oxidative damage, these findings also raise intriguing questions about the physiological role of MAGEA genes in maintaining germ cell metabolic and redox homeostasis. Collectively, our findings highlight how understanding the molecular functions of MAGEA3/6 in both germ cells and cancer may uncover previously unrecognized metabolic vulnerabilities and potential therapeutic opportunities in pancreatic cancer.

## Author Contributions

Conceptualization, S.T., V.J., and K.F.T.; methodology, S.T., V.J., B.B.V., J.S.S.G., R.R.F.G., D.Š., M.C.H.S., S.U., J.K., L.S., E.B., J.P., M.J., M.V., P.R.P., and K.F.T.; validation, S.T. and V.J.; data analysis, S.T., D.Š., J.S.S.G., M.C.H.S., and R.R.F.G.; investigation, S.T., V.J., and B.B.V.; resources, E.B., P.R.P., and K.F.T.; data curation, S.T., V.J., J.S.S.G., M.C.H.S., D.Š., and K.F.T.; writing—original draft preparation, S.T. and K.F.T.; writing—review and editing, S.T., V.J., B.B.V., and K.F.T.; visualization, S.T., D.Š., and K.F.T.; supervision, K.F.T.; project administration, K.F.T.; funding acquisition, K.F.T.

## FUNDING

Funding for this work was supported by American Cancer Society Research Scholar Award (181691010 to PRP); and Cancer Prevention and Research Institute of Texas (R1117 to PRP, RR200059 to KFT); the Texas Tech University start-up and the Texas Center for Comparative Cancer Research (TC3R) (KFT); Foundation for Prader–Willi Syndrome Research Grants (22-0321 and 23-0447 to KFT); Fulbright fellowship (to BBV); ASEF Global Fellowship (to SU, VJ, JK), Slovenian Research and Innovation Agency (ARIS) grants (P1-0245, J7-70250, BI-US/22-24-007, BI-US/24-26-022) and the American Lebanese Syrian Associated Charities of St Jude Children’s Research Hospital.

## DATA AVAILABILITY STATEMENT

The raw data supporting the conclusions of this article will be made available by the corresponding author upon reasonable request. The raw and processed RNAseq data are available in the Gene Expression Omnibus database from the NCBI under accession number XXXXXX.

## ACKNOWLEDGEMENTS

We thank the members of the Fon Tacer laboratory for their valuable advice and critical discussions. We also acknowledge the Texas Tech University School of Veterinary Medicine, the Department of Biological Sciences, the Texas Tech University Graduate School, and the Texas Tech University Association of Biologists for support through Grants-in-Aid. The results presented in this study are partly based on data generated by the Cancer Cell Line Encyclopedia (CCLE).

## CONFLICTS OF INTEREST

The authors declare no conflict of interest.

---

**Figure S1.**
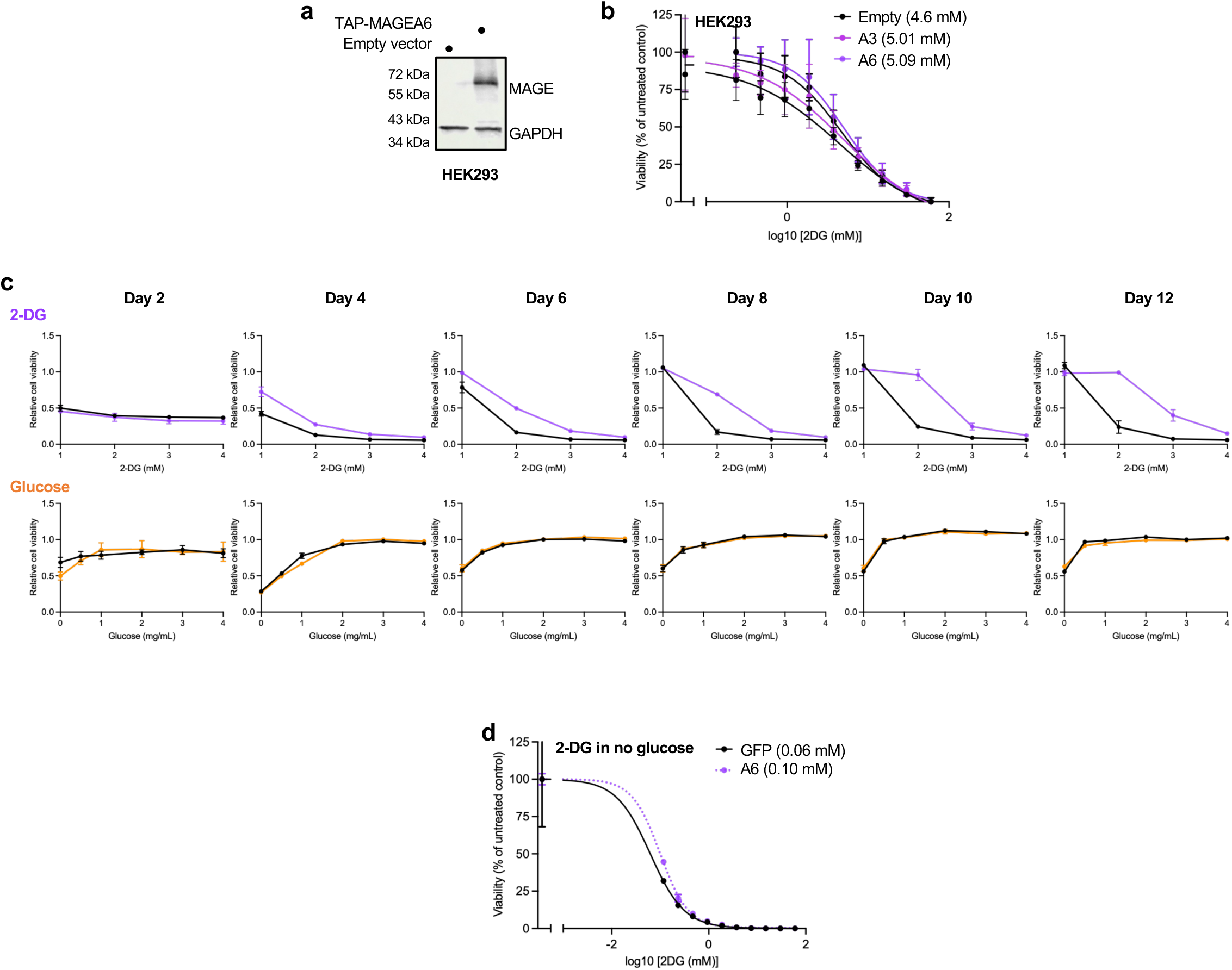
MAGEA6 promotes resistance to 2-deoxyglucose (2-DG)-induced metabolic stress in pancreatic cancer cells. **a.** Western blot analysis confirming MAGEA6 expression in HEK293 cells. **b.** Dose–response curves showing the sensitivity of HEK293 cells expressing empty vector, MAGEA3, or MAGEA6 to 2-DG, assessed using the Alamar Blue assay five days after treatment. Cell viability is expressed as a percentage of the untreated control, with IC₅₀ values indicated in mM. **c.** Longitudinal analysis of GFP-control and MAGEA6-expressing MIA PaCa-2 cells exposed to increasing concentrations of 2-DG (top; purple) or cultured in media containing varying glucose concentrations (bottom; orange). Cell viability was assessed using the Alamar Blue assay and normalized to day 0 (baseline). **d.** Dose-response curves showing the sensitivity of GFP-control and MAGEA6-expressing MIA PaCa-2 cells to 2-DG under glucose-free culture conditions. Cell viability is expressed as a percentage of the untreated control, with IC₅₀ values indicated in mM. Data are presented as mean ± SD of four technical replicates. Experiments were independently repeated at least three times, and one representative experiment is shown.

**Figure S2.**
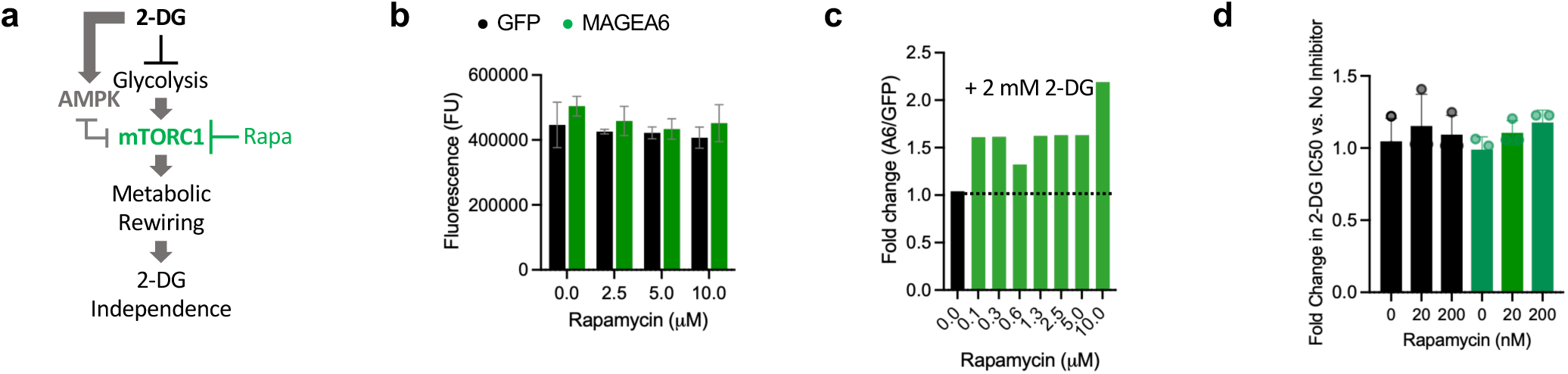
MAGEA6-mediated resistance to 2-DG is largely independent of AMPK–mTOR signaling in pancreatic cancer cells. **a.** Schematic representation of glycolytic inhibition by 2-DG and the AMPK–mTOR signaling axis, highlighting its downstream regulation of cellular growth and metabolism. **b.** Cell viability of GFP-control and MAGEA6-expressing MIA PaCa-2 cells following 7 days of treatment with increasing concentrations of rapamycin, assessed using the Alamar Blue assay and expressed as fluorescence units. **c.** Fold change in cell viability of MAGEA6-expressing cells relative to GFP controls following 5 days of combined rapamycin and 2 mM 2-DG treatment. **d.** Fold change in 2-DG IC₅₀ values following treatment with 20 or 200 nM rapamycin relative to the corresponding controls without rapamycin. Data are presented as mean ± SD. Statistical significance was assessed by one-way ANOVA followed by Šidák’s multiple-comparisons test, as indicated. \*\*\**P* < 0.0001.

**Figure S3.**
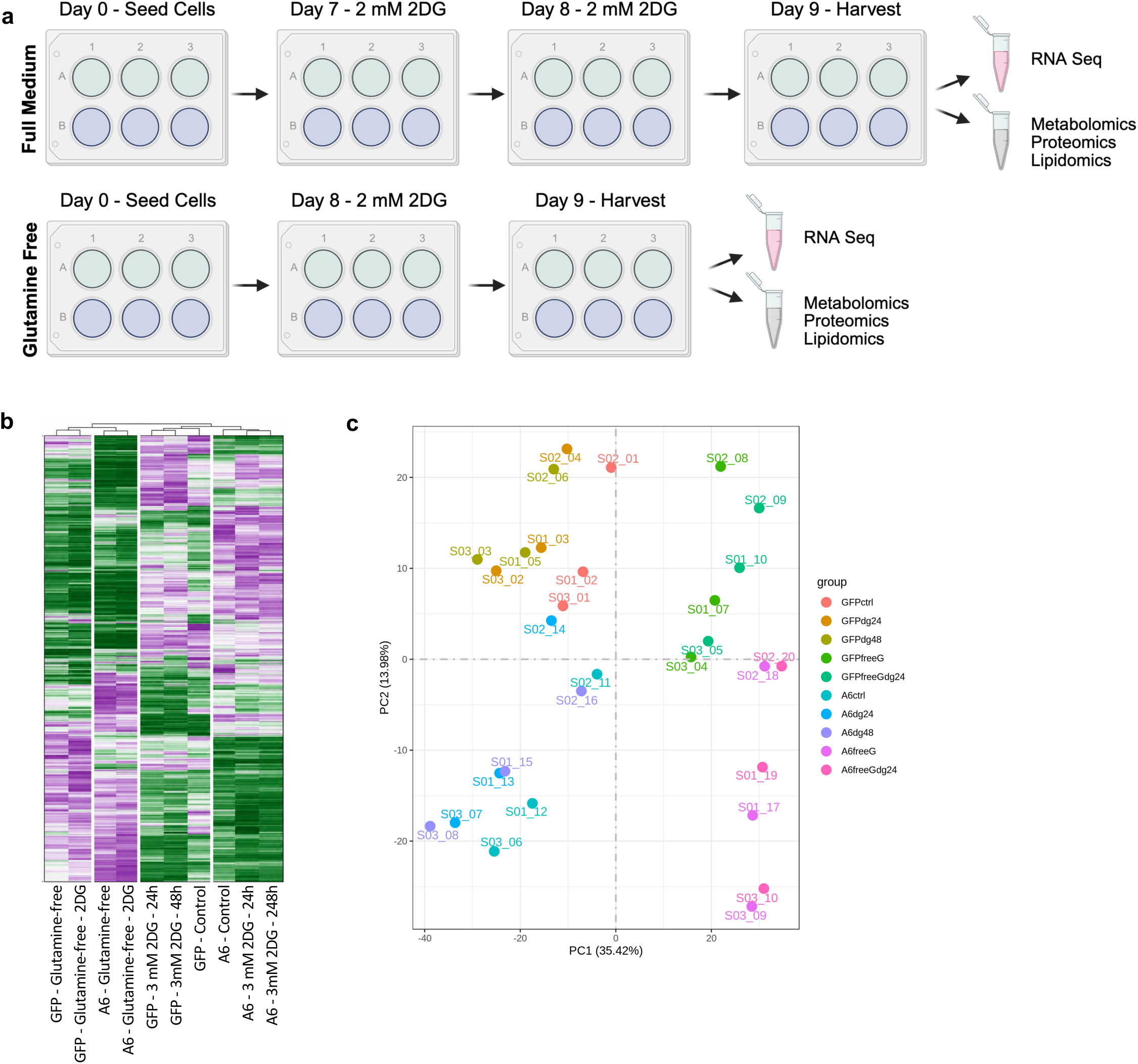
Multi-omics analysis reveals distinct molecular profiles associated with MAGEA6 expression and metabolic stress in pancreatic cancer cells. **a.** Schematic representation of the experimental design for integrated transcriptomic, proteomic, metabolomic, and lipidomic analyses of GFP-control and MAGEA6-expressing MIA PaCa-2 cells under different metabolic conditions. Cells were cultured in complete or glutamine-free medium and treated with 2 mM 2-DG for 24 or 48 h, as indicated. Samples were collected for RNA sequencing (RNA-seq), proteomics, metabolomics, and lipidomics, with three biological replicates per experimental condition. The analyses presented in this figure focus on RNA-seq data. **b.** Hierarchical clustering heatmap of RNA-seq gene expression profiles in GFP-control and MAGEA6-expressing MIA PaCa-2 cells under the indicated metabolic conditions. Rows represent genes, and columns represent experimental groups. Relative gene expression is displayed using a green–white–purple color scale, with green indicating lower expression, white indicating intermediate expression, and purple indicating higher expression. Hierarchical clustering reveals distinct transcriptional patterns associated with MAGEA6 expression, 2-DG treatment, and glutamine deprivation. **c.** Principal component analysis (PCA) of RNA-seq data from GFP-control and MAGEA6-expressing MIA PaCa-2 cells under the indicated metabolic conditions. Each point represents an individual biological replicate, colored according to genotype and treatment condition. Principal components 1 (PC1) and 2 (PC2) explain 35.42% and 13.98% of the total variance, respectively. The distribution of samples illustrates differences in global transcriptional profiles associated with MAGEA6 expression and metabolic stress.

**Figure S4.**
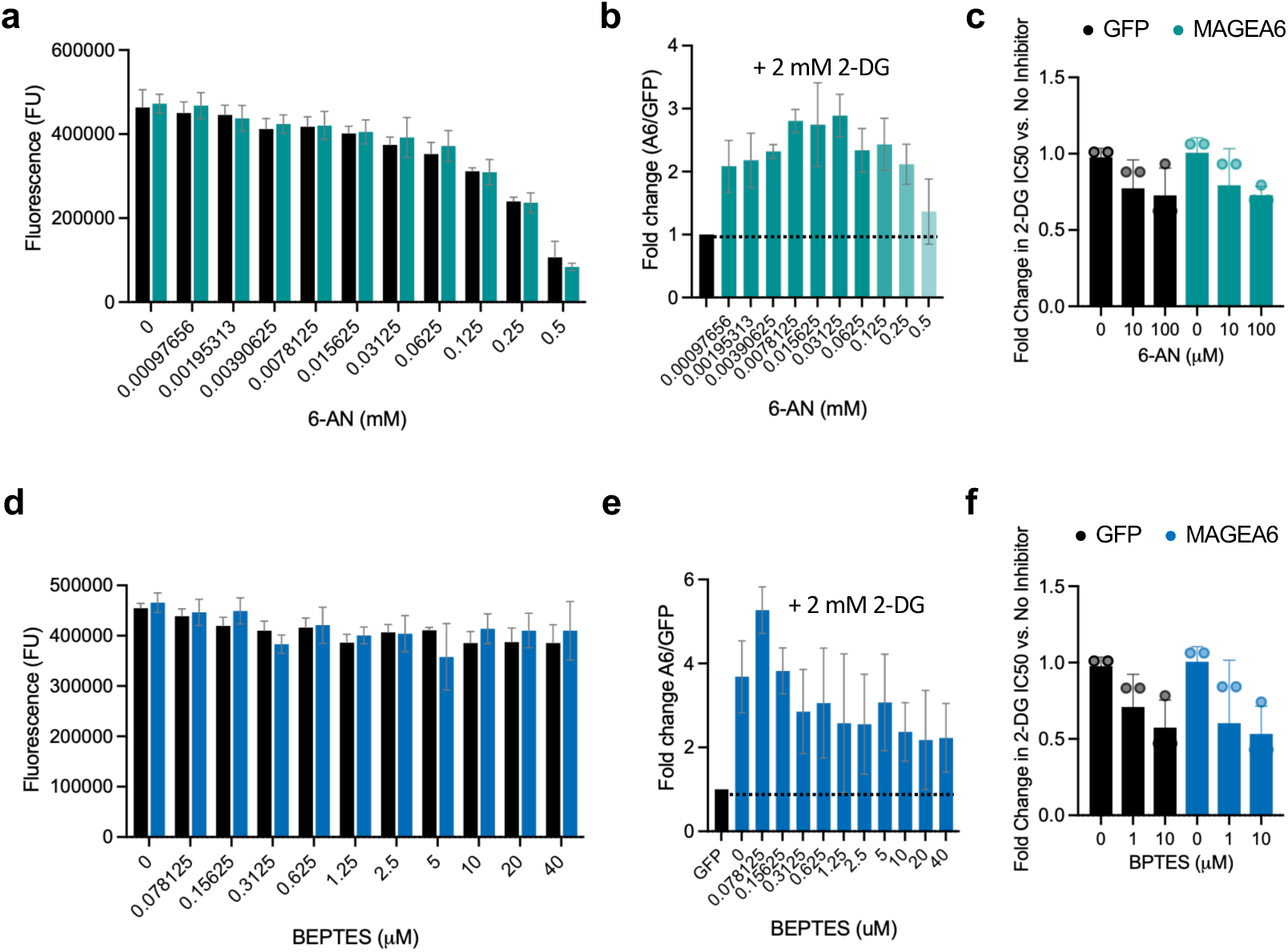
Effects of pentose phosphate pathway and glutaminase inhibition on MAGEA6-associated resistance to 2-DG in MIA PaCa-2 pancreatic cancer cells. **a.** Viability of GFP-control and MAGEA6-expressing MIA PaCa-2 cells following treatment with increasing concentrations of the pentose phosphate pathway (PPP) inhibitor 6-aminonicotinamide (6-AN), assessed using the Alamar Blue assay and expressed as fluorescence units (FU). **b.** Fold change in cell viability of MAGEA6-expressing cells relative to GFP controls following treatment with increasing concentrations of 6-AN in combination with 2 mM 2-DG. **c.** Fold change in 2-DG half-maximal inhibitory concentration (IC50) values following treatment with the indicated concentrations of 6-AN relative to the corresponding controls without 6-AN. **d.** Viability of GFP-control and MAGEA6-expressing MIA PaCa-2 cells following treatment with increasing concentrations of the glutaminase inhibitor BPTES, assessed using the Alamar Blue assay and expressed as fluorescence units (FU). **e.** Fold change in cell viability of MAGEA6-expressing cells relative to GFP controls following treatment with increasing concentrations of BPTES in combination with 2 mM 2-DG. **f.** Fold change in 2-DG IC₅₀ values following treatment with the indicated concentrations of BPTES relative to the corresponding controls without BPTES. Data are presented as mean ± SD. Statistical significance was assessed only for panels c and f using one-way ANOVA followed by Šidák’s multiple-comparisons test.

**Figure S5.**
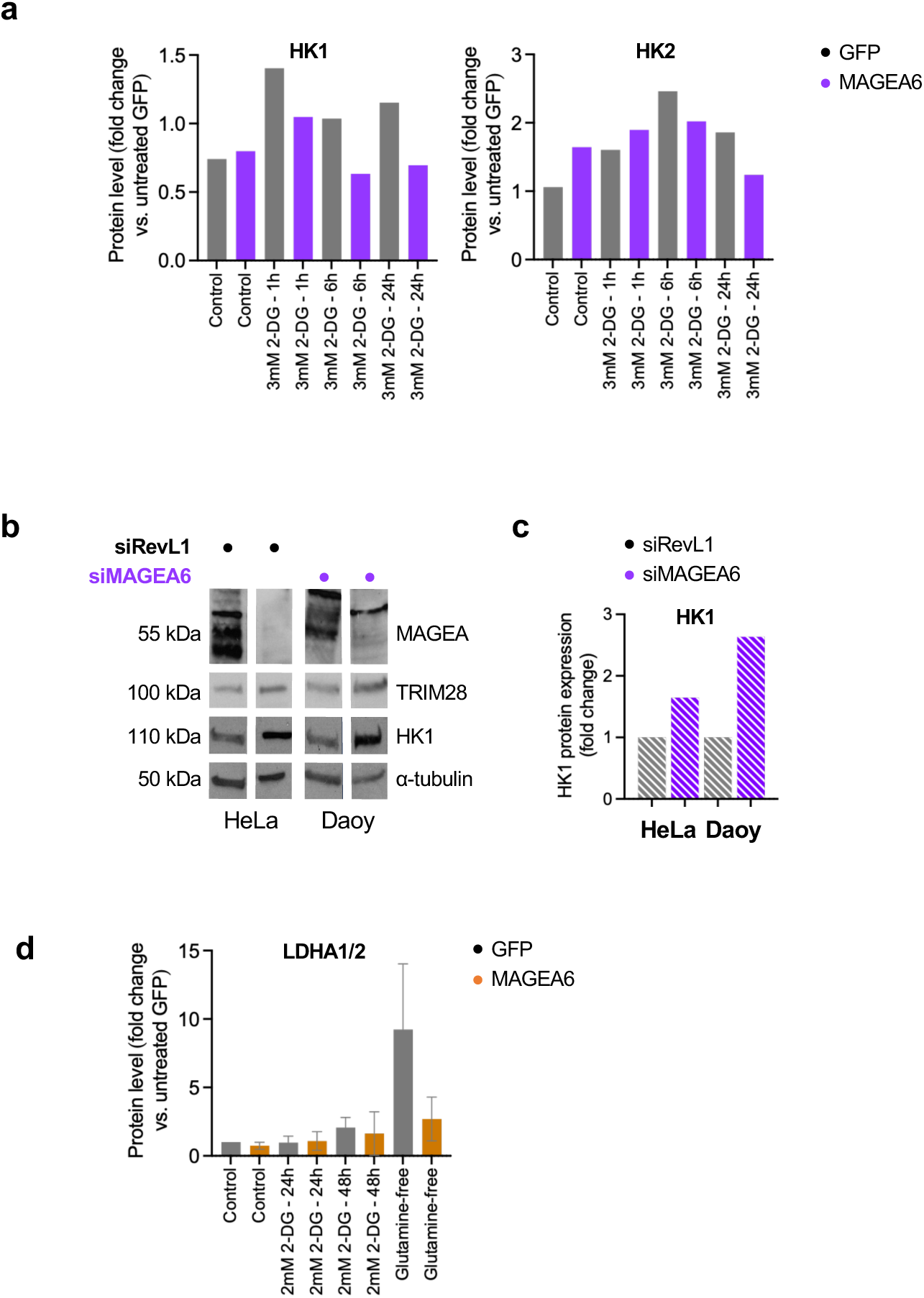
MAGEA6 differentially regulates glycolytic enzyme abundance under metabolic stress and is associated with hexokinases. **a.** Densitometric quantification of HK1 and HK2 protein abundance normalized to TRIM28 and expressed as fold change relative to untreated GFP-control cells, based on the Western blots shown in Figure 5b. **b.** Representative Western blots showing MAGEA6, HK1, and TRIM28 protein abundance following MAGEA6 silencing in HeLa and Daoy cells. α-Tubulin was used as an additional loading control. **c.** Densitometric quantification of HK1 protein abundance in Daoy cells following MAGEA6 silencing, normalized to the corresponding loading control and expressed as fold change relative to control cells. **d.** Densitometric quantification of the combined LDHA1/2 bands, expressed as fold change relative to untreated GFP-control cells, based on the Western blots shown in Figure 5e. Data are presented as mean ± SD, where applicable.

## Notes

### Competing Interest Statement

The authors have declared no competing interest.

## REFERENCES

1. Mannucci, A. and A. Goel, Advances in pancreatic cancer early diagnosis, prevention, and treatment: The past, the present, and the future. CA: A Cancer Journal for Clinicians, 2025.

2. Robatel, S. and M. Schenk, Current limitations and novel perspectives in pancreatic cancer treatment. Cancers, 2022. 14(4): p. 985.

3. Halbrook, C.J., et al., Pancreatic cancer: advances and challenges. Cell, 2023. 186(8): p. 1729–1754.

4. Society, A.C. American Cancer Society’s Cancer Statistics. 2025; Available from: https://www.cancer.org/cancer/types/pancreatic-cancer/detection-diagnosis-staging/survival-rates.html.

5. Yu, B., S. Shao, and W. Ma, Frontiers in pancreatic cancer on biomarkers, microenvironment, and immunotherapy. Cancer letters, 2025. 610: p. 217350.

6. .

7. O’Reilly, E.M., et al., Daraxonrasib or Chemotherapy in Previously Treated Metastatic Pancreatic Cancer. N Engl J Med, 2026. 395(4): p. 325–337.

8. Mannucci, A. and A. Goel, Advances in pancreatic cancer early diagnosis, prevention, and treatment: The past, the present, and the future. CA Cancer J Clin, 2026. 76(1): p. e70035.

9. Giurini, E.F., et al., Looking Beyond Checkpoint Inhibitor Monotherapy: Uncovering New Frontiers for Pancreatic Cancer Immunotherapy. Mol Cancer Ther, 2025. 24(1): p. 18–32.

10. Hu, Z.I. and E.M. O’Reilly, Therapeutic developments in pancreatic cancer. Nature Reviews Gastroenterology & Hepatology, 2024. 21(1): p. 7–24.

11. Bayat, T., et al., Cross-Species Upregulation of MAGED2 in Liver Cancer Suggests a Role in Obesity-Driven Tumor Progression. Current Issues in Molecular Biology, 2026. 48(2): p. 139.

12. Collins, R.R.J., et al., Melanoma antigens in pediatric medulloblastoma contribute to tumor heterogeneity and species-specificity of group 3 tumors. Acta Neuropathol Commun, 2025. 13(1): p. 164.

13. Florke Gee, R.R., et al., Emerging roles of the MAGE protein family in stress response pathways. J Biol Chem, 2020. 295(47): p. 16121–16155.

14. Fon Tacer, K., et al., MAGE cancer-testis antigens protect the mammalian germline under environmental stress. Science Advances, 2019. 5(5): p. eaav4832.

15. Lee, A.K. and P.R. Potts, A Comprehensive Guide to the MAGE Family of Ubiquitin Ligases. J Mol Biol, 2017. 429(8): p. 1114–1142.

16. Weon, J.L. and P.R. Potts, The MAGE protein family and cancer. Current opinion in cell biology, 2015. 37: p. 1–8.

17. van der Bruggen, P., et al., A gene encoding an antigen recognized by cytolytic T lymphocytes on a human melanoma. Science, 1991. 254(5038): p. 1643-7.

18. Barnett, K.K., et al., FDA Approval Summary: Afamitresgene Autoleucel for Adults with HLA-Restricted, MAGE-A4-Positive Unresectable or Metastatic Synovial Sarcoma after Prior Chemotherapy. Clin Cancer Res, 2025. 31(15): p. 3112–3117.

19. Qin, H., et al., Immunization with a multi-antigen targeted DNA vaccine eliminates chemoresistant pancreatic cancer by disrupting tumor-stromal cell crosstalk. Journal of Translational Medicine, 2023. 21(1): p. 702.

20. Das, B. and S. Senapati, Functional and mechanistic studies reveal MAGEA3 as a pro-survival factor in pancreatic cancer cells. Journal of Experimental & Clinical Cancer Research, 2019. 38(1): p. 294.

21. Kim, J., et al., The clinical significance of MAGEA3 expression in pancreatic cancer. International journal of cancer, 2006. 118(9): p. 2269–2275.

22. Ravichandran, R., et al., Regulation of MAGE-A3/6 by the CRL 4-DCAF 12 ubiquitin ligase and nutrient availability. EMBO reports, 2019. 20(7): p. e47352.

23. Su, J., et al., Unveiling MAGEA3: a novel predictive biomarker for bevacizumab resistance in colorectal cancer. Cancer Drug Resistance, 2025. 8: p. 22.

24. Su, J., et al., Unveiling MAGEA3: a novel predictive biomarker for bevacizumab resistance in colorectal cancer. Cancer Drug Resist, 2025. 8: p. 22.

25. Gan, Y., et al., Cancer testis antigen MAGEA3 in serum and serum-derived exosomes serves as a promising biomarker in lung adenocarcinoma. Scientific Reports, 2024. 14(1): p. 7573.

26. Poojary, M., P.V. Jishnu, and S.P. Kabekkodu, Prognostic value of melanoma-associated antigen-A (MAGE-A) gene expression in various human cancers: a systematic review and meta-analysis of 7428 patients and 44 studies. Molecular Diagnosis & Therapy, 2020. 24(5): p. 537–555.

27. Schaefer, P., T. Paraschiakos, and S. Windhorst, Oncogenic activity and cellular functionality of melanoma associated antigen A3. Biochemical pharmacology, 2021. 192: p. 114700.

28. Fon Tacer, K., et al., MAGE cancer-testis antigens protect the mammalian germline under environmental stress. Sci Adv, 2019. 5(5): p. eaav4832.

29. Griswold, M.D., Spermatogenesis: The Commitment to Meiosis. Physiol Rev, 2016. 96(1): p. 1–17.

30. Doyle, J.M., et al., MAGE-RING protein complexes comprise a family of E3 ubiquitin ligases. Molecular cell, 2010. 39(6): p. 963–974.

31. Jin, X., et al., MAGE-TRIM28 complex promotes the Warburg effect and hepatocellular carcinoma progression by targeting FBP1 for degradation. Oncogenesis, 2017. 6(4): p. e312–e312.

32. Monte, M., et al., MAGE-A tumor antigens target p53 transactivation function through histone deacetylase recruitment and confer resistance to chemotherapeutic agents. Proceedings of the National Academy of Sciences, 2006. 103(30): p. 11160–11165.

33. Pineda, C.T., et al., Degradation of AMPK by a cancer-specific ubiquitin ligase. Cell, 2015. 160(4): p. 715–728.

34. Yang, B., et al., MAGE-A, mMage-b, and MAGE-C proteins form complexes with KAP1 and suppress p53-dependent apoptosis in MAGE-positive cell lines. Cancer research, 2007. 67(20): p. 9954–9962.

35. Jones, R.G., et al., AMP-activated protein kinase induces a p53-dependent metabolic checkpoint. Molecular cell, 2005. 18(3): p. 283–293.

36. Shukla, S.A., et al., Cancer-germline antigen expression discriminates clinical outcome to CTLA-4 blockade. Cell, 2018. 173(3): p. 624–633. e8.

37. Tsang, Y.H., et al., Differential expression of MAGEA6 toggles autophagy to promote pancreatic cancer progression. Elife, 2020. 9: p. e48963.

38. Pan, S.-J., et al., MAGEA6 promotes human glioma cell survival via targeting AMPKα1. Cancer letters, 2018. 412: p. 21–29.

39. Shaw, R.J., Tumor metabolism: MAGE-A proteins help TRIM turn over AMPK. Current Biology, 2015. 25(10): p. R418–R420.

40. Wu, F., et al., miR-1273g silences MAGEA3/6 to inhibit human colorectal cancer cell growth via activation of AMPK signaling. Cancer letters, 2018. 435: p. 1-9.

41. Ye, X., et al., Knockdown of MAGEA6 activates AMP-activated protein kinase (AMPK) signaling to inhibit human renal cell carcinoma cells. Cellular Physiology and Biochemistry, 2018. 45(3): p. 1205–1218.

42. Dong, C., et al., Loss of FBP1 by Snail-mediated repression provides metabolic advantages in basal-like breast cancer. Cancer cell, 2013. 23(3): p. 316–331.

43. Biancur, D.E., et al., Functional Genomics Identifies Metabolic Vulnerabilities in Pancreatic Cancer. Cell Metab, 2021. 33(1): p. 199–210.e8.

44. Kamphorst, J.J., et al., Hypoxic and Ras-transformed cells support growth by scavenging unsaturated fatty acids from lysophospholipids. Proc Natl Acad Sci U S A, 2013. 110(22): p. 8882–7.

45. Qin, C., et al., Metabolism of pancreatic cancer: paving the way to better anticancer strategies. Molecular Cancer, 2020. 19(1): p. 50.

46. 46. Costamagna, A., et al., Lipid signaling networks in pancreatic cancer progression and therapeutic perspectives. Trends in Endocrinology & Metabolism, 2026.

47. Coleman, M.C., et al., 2-Deoxy-d-glucose causes cytotoxicity, oxidative stress, and radiosensitization in pancreatic cancer. Free Radical Biology and Medicine, 2008. 44(3): p. 322–331.

48. Pusapati, R.V., et al., mTORC1-Dependent Metabolic Reprogramming Underlies Escape from Glycolysis Addiction in Cancer Cells. Cancer Cell, 2016. 29(4): p. 548–562.

49. Sasaki, K., et al., Nanoparticle-Mediated Delivery of 2-Deoxy-D-Glucose Induces Antitumor Immunity and Cytotoxicity in Liver Tumors in Mice. Cellular and Molecular Gastroenterology and Hepatology, 2021. 11(3): p. 739–762.

50. Xie, B.W., et al., Dual-wavelength imaging of tumor progression by activatable and targeting near-infrared fluorescent probes in a bioluminescent breast cancer model. PLoS One, 2012. 7(2): p. e31875.

51. Kim, D., et al., Graph-based genome alignment and genotyping with HISAT2 and HISAT-genotype. Nature Biotechnology, 2019. 37(8): p. 907–915.

52. Love, M.I., W. Huber, and S. Anders, Moderated estimation of fold change and dispersion for RNA-seq data with DESeq2. Genome Biology, 2014. 15(12): p. 550.

53. Bustin, S.A., et al., The MIQE Guidelines: Minimum Information for Publication of Quantitative Real-Time PCR Experiments. Clinical Chemistry, 2009. 55(4): p. 611–622.

54. Schindelin, J., et al., Fiji: an open-source platform for biological-image analysis. Nature Methods, 2012. 9(7): p. 676–682.

55. Kennedy, G.T., et al., Targeted Intraoperative Molecular Imaging for Localizing Nonpalpable Tumors and Quantifying Resection Margin Distances. JAMA Surg, 2021. 156(11): p. 1043–1050.

56. Chhabra, R. and M. Nanjundan, Lysophosphatidic acid reverses Temsirolimus-induced changes in lipid droplets and mitochondrial networks in renal cancer cells. PLoS One, 2020. 15(6): p. e0233887.

57. Minami, J.K., et al., CDKN2A deletion remodels lipid metabolism to prime glioblastoma for ferroptosis. Cancer Cell, 2023. 41(6): p. 1048–1060.e9.

58. Bogataj, U., et al., Heterogenous mitochondrial ultrastructure and metabolism of human glioblastoma cells: differences between stem-like and differentiated cancer cells in response to chemotherapy. Radiol Oncol, 2025. 59(4): p. 551–565.

59. Guzmán, C., et al., ColonyArea: an ImageJ plugin to automatically quantify colony formation in clonogenic assays. PLoS One, 2014. 9(3): p. e92444.

60. Doğru, D., et al., An automated in vitro wound healing microscopy image analysis approach utilizing U-net-based deep learning methodology. BMC Med Imaging, 2024. 24(1): p. 158.

61. Molefe, P.F., et al., The In Vitro Wound-Scratch Assay: Applications, Technical Advances, and Limitations in Wound Healing Research. International Wound Journal, 2026. 23(6): p. e70964.

62. Justus, C.R., et al., In vitro cell migration and invasion assays. J Vis Exp, 2014(88).

63. Zhang, Y., et al., The Glutaminase-1 Inhibitor [(11)C-carbony]BPTES: Synthesis and Positron Emission Tomography Study in Mice. Pharmaceuticals (Basel), 2023. 16(7).

64. Niepel, M., et al., Measuring Cancer Drug Sensitivity and Resistance in Cultured Cells. Curr Protoc Chem Biol, 2017. 9(2): p. 55–74.

65. Shoshan, M.C., 3-bromopyruvate: Targets and outcomes. Journal of Bioenergetics and Biomembranes, 2012. 44(1): p. 7–15.

66. Stuelten, C.H., C.A. Parent, and D.J. Montell, Cell motility in cancer invasion and metastasis: insights from simple model organisms. Nat Rev Cancer, 2018. 18(5): p. 296–312.

67. Pelicano, H., et al., Glycolysis inhibition for anticancer treatment. Oncogene, 2006. 25(34): p. 4633–4646.

68. Zhang, D., et al., 2-Deoxy-D-glucose targeting of glucose metabolism in cancer cells as a potential therapy. Cancer Letters, 2014. 355(2): p. 176–183.

69. Ye, X.-T., et al., Knockdown of MAGEA6 Activates AMP-Activated Protein Kinase (AMPK) Signaling to Inhibit Human Renal Cell Carcinoma Cells. Cellular Physiology and Biochemistry, 2018. 45: p. 1205–1218.

70. Zhou, J., et al., Knockdown of MAGE-A6 enhanced the irradiation sensitivity of non-small cell lung cancer cells by activating the AMPK pathway. Environmental Toxicology, 2022. 37(7): p. 1711–1722.

71. Zhu, H., et al., Targeting oncogenic MAGEA6 sensitizes triple negative breast cancer to doxorubicin through its autophagy and ferroptosis by stabling AMPKα1. Cell Death Discovery, 2024. 10(1): p. 430.

72. Mandic, M., et al., Multifaceted role of AMPK in autophagy: more than a simple trigger? American Journal of Physiology-Cell Physiology, 2025. 329(5): p. C1380–C1397.

73. Daneshmandi, S., et al., 6-Phosphogluconate dehydrogenase promotes mitochondrial fusion and immune suppression in tumor-associated monocytic suppressor cells. Nature Communications, 2026. 17(1): p. 229.

74. Son, J., et al., Glutamine supports pancreatic cancer growth through a KRAS-regulated metabolic pathway. Nature, 2013. 496(7443): p. 101–105.

75. Ravichandran, R., et al., Regulation of MAGE-A3/6 by the CRL4-DCAF12 ubiquitin ligase and nutrient availability. EMBO Reports, 2019. 20(7): p. EMBR201847352.

76. Lohman, D.C., et al., Mitochondrial COQ9 is a lipid-binding protein that associates with COQ7 to enable coenzyme Q biosynthesis. Proceedings of the National Academy of Sciences, 2014. 111(44): p. E4697–E4705.

77. Srivastava, S., et al., Regulation of mitochondrial protein import by the nucleotide exchange factors GrpEL1 and GrpEL2 in human cells. Journal of Biological Chemistry, 2017. 292(44): p. 18075–18090.

78. Stroud, D.A., et al., Accessory subunits are integral for assembly and function of human mitochondrial complex I. Nature, 2016. 538(7623): p. 123–126.

79. Timón-Gómez, A., et al., Distinct Roles of Mitochondrial HIGD1A and HIGD2A in Respiratory Complex and Supercomplex Biogenesis. Cell Reports, 2020. 31(5).

80. Eckert, R.L., Transglutaminase 2 takes center stage as a cancer cell survival factor and therapy target. Mol Carcinog, 2019. 58(6): p. 837–853.

81. McMaster, C.R., From yeast to humans - roles of the Kennedy pathway for phosphatidylcholine synthesis. FEBS Lett, 2018. 592(8): p. 1256–1272.

82. Porporato, P.E., et al., Mitochondrial metabolism and cancer. Cell Research, 2018. 28(3): p. 265–280.

83. Cheng, G., et al., Profiling and targeting of cellular bioenergetics: inhibition of pancreatic cancer cell proliferation. British Journal of Cancer, 2014. 111(1): p. 85–93.

84. Yan, H., et al., Dual Metabolic Blockade in Pancreatic Cancer: Potent Anticancer Activity of Mitochondria-Targeted Glycolysis and OXPHOS Inhibitors. Journal of Medicinal Chemistry, 2026. 69(11): p. 13452–13472.

85. Cheng, Y., et al., High glucose-induced resistance to 5-fluorouracil in pancreatic cancer cells alleviated by 2-deoxy-D-glucose. Biomed Rep, 2014. 2(2): p. 188–192.

86. Dai, S., et al., Glycolysis promotes the progression of pancreatic cancer and reduces cancer cell sensitivity to gemcitabine. Biomedicine & Pharmacotherapy, 2020. 121: p. 109521.

87. Goldberg, L., R. Israeli, and Y. Kloog, FTS and 2-DG induce pancreatic cancer cell death and tumor shrinkage in mice. Cell Death & Disease, 2012. 3(3): p. e284–e284.

88. Malm, S.W., et al., The anti-tumor efficacy of 2-deoxyglucose and D-allose are enhanced with p38 inhibition in pancreatic and ovarian cell lines. Journal of Experimental & Clinical Cancer Research, 2015. 34(1): p. 31.

89. Shukla, H.D., et al., 3-Bromopyruvate in Combination with Radiation Inhibits Pancreatic Tumor Growth by Suppressing Hexokinase II (HK2) Enzyme of Aerobic Glycolysis. International Journal of Radiation Oncology, Biology, Physics, 2025. 123(1): p. e509–e510.

90. Xiao, H., et al., Separate and concurrent use of 2-deoxy-D-glucose and 3-bromopyruvate in pancreatic cancer cells. Oncol Rep, 2013. 29(1): p. 329–34.

91. Shimoni-Sebag, A., et al., A metabolic switch to the pentose-phosphate pathway induces radiation resistance in pancreatic cancer. Radiotherapy and Oncology, 2025. 202: p. 110606.

92. Pineda, C.T. and P.R. Potts, Oncogenic MAGEA-TRIM28 ubiquitin ligase downregulates autophagy by ubiquitinating and degrading AMPK in cancer. Autophagy, 2015. 11(5): p. 844–846.

93. Pesce, N.A., et al., HK1 and HK2 Beyond Glycolysis: Mitochondrial Interactions and Dual Roles in Metabolism and Cell Fate. Adv Biol (Weinh), 2026. 10(1): p. e00472.

94. Jiang, M., et al., A pan-cancer analysis of molecular characteristics and oncogenic role of hexokinase family genes in human tumors. Life Sciences, 2021. 264: p. 118669.

95. Ravichandran, R., et al., Regulation of MAGE-A3/6 by the CRL4-DCAF12 ubiquitin ligase and nutrient availability. EMBO Rep, 2019. 20(7): p. e47352.

96. Viale, A., et al., Oncogene ablation-resistant pancreatic cancer cells depend on mitochondrial function. Nature, 2014. 514(7524): p. 628–632.

97. Ding, Y., et al., Glutamine Metabolism in Health and Diseases. MedComm, 2026. 7(10): p. e71035.

98. Jin, J., et al., Targeting glutamine metabolism as a therapeutic strategy for cancer. Experimental & Molecular Medicine, 2023. 55(4): p. 706–715.

99. Krahmer, N., et al., Phosphatidylcholine Synthesis for Lipid Droplet Expansion Is Mediated by Localized Activation of CTP:Phosphocholine Cytidylyltransferase. Cell Metabolism, 2011. 14(4): p. 504–515.

